# Evidence for vocal signatures and voice-prints in a wild parrot

**DOI:** 10.1101/2023.01.20.524864

**Authors:** Simeon Q. Smeele, Juan Carlos Senar, Lucy M. Aplin, Mary Brooke McElreath

**Author notes:** **Author for correspondence:** Simeon Q. Smeele. Joint senior author (Lucy M. Aplin, Mary Brooke McElreath).

## Abstract

In humans, identity is partly encoded in a voice-print that is carried across multiple vocalisations. Other species also signal vocal identity in calls, such as shown in the contact call of parrots. However, it remains unclear to what extent other call types in parrots are individually distinct, and whether there is an analogous voice-print across calls. Here we test if an individual signature is present in other call types, how stable this signature is, and if parrots exhibit voice-prints across call types. We recorded 5599 vocalisations from 229 individually-marked monk parakeets (*Myiopsitta monachus*) over a two year period in Barcelona, Spain. We examined five distinct call types, finding evidence for an individual signature in three. We further show that in the contact call, while birds are individually distinct, the calls are more variable than previously assumed, changing over short time scales (seconds to minutes). Finally, we provide evidence for voice-prints across multiple call types, with a discriminant function being able to predict caller identity across call types. This suggests that monk parakeets may be able to use vocal cues to recognise conspecifics, even across vocalisation types and without necessarily needing active vocal signatures of identity.

## Introduction

Individual recognition and signalling of individual identity can play an important role in social interactions and decision-making. Examples of how individuals can benefit from individual recognition are wide-ranging, and include helping relatives (Russell and Hatchwell 2001), remembering reliable cooperators (Boesch 1994) and strategically directing aggression (Hobson, Mønster, and DeDeo 2021). For the individual that is recognised, signalling identity is beneficial if the benefits associated with incurring affiliative behaviour outweigh potential costs associated with misidentification (Johnstone 1997). While it sometimes also pays to hide identity (Tibbetts and Dale 2007; Johnstone 1997; Carlson, Kelly, and Couzin 2020), in most cases, the benefits of broadcasting identity likely outweigh the potential costs. In fission-fusion societies, for instance, signalling identity may allow individuals to preferentially reassociate with a subset of the population when confronted with a large number of potential interaction partners (Kummer 2017; Aureli, Schaffner, and Schino 2022). Early human societies were fission-fusion based and likely heavily dependent on cooperation between individuals (Migliano and Vinicius 2022); perhaps not surprisingly, the human face has evolved to allow for maximum individual distinctiveness (Sheehan and Nachman 2014).

Across species, individual identity has been found to be conveyed through multiple potential sensory modalities, including olfactory, acoustic or visual cues. For example, several social wasps display distinctive facial features (Tibbetts 2004). However, while visual or olfactory distinctiveness is useful during close interactions, they are likely less effective across longer distances or in low visibility environments such as tropical forests or turbid waters. Vocal signals are much better suited for these situations, and vocal broadcasting of identity has been found across a wide range of taxonomic groups, ranging from American Goldfinches (*Spinus tristis*) (Mundinger 1970) to bottle-nosed dolphins (*Tursiops truncatus*) (Janik and Sayigh 2013). These species often have one call type that is very stereotyped within individuals, with enough structural complexity to allow for many unique variants. For example, bottle-nosed dolphins produce a very stereotyped signature whistle when out of visual contact, where the individual signature is encoded in the frequency modulation pattern, or in other words how the frequency goes up and down (Janik, Sayigh, and Wells 2006). Individuals predominantly produce ‘their’ signature whistle, and the duration combined with the frequency modulation allows for many unique patterns.

While a single vocal signal to broadcast identity is useful, individuals will often produce multiple call types, and could therefore benefit from being recognised across these calls. Three potential solutions to the need to be recognised in multiple call types are possible (see Figure 1). The first is making each call type individually distinct. Such a strategy has been shown in a variety of bird species (Elie and Theunissen 2018; Charrier et al. 2001; Mä kelin et al. 2021), bats (Prat, Taub, and Yovel 2016) and some primate species (Keenan et al. 2020; Bouchet et al. 2012; Salmi, Hammerschmidt, and Doran-Sheehy 2014). However, maintaining multiple signals of identity is cognitively demanding for signallers and receivers to remember; consequently, this strategy is likely constrained to species with either small vocal repertoires or small group sizes (Elie and Theunissen 2018). The second solution is to combine a single identity call with the other call types in a sequence (Rauber, Kranstauber, and Manser 2020). The cognitive demands of this strategy are much lower, and if flexibly deployed, it potentially allows individuals to signal identity in contexts where recognition is beneficial and hide identity in other contexts. However, it increases the complexity and potential cost of vocal production, as all individually distinct vocalisations now involve at least two elements. The third solution is to evolve a recognisable voice-print across call types. This can be achieved via the specific morphology of the vocal production organ, leaving a unique and recognisable cue on all vocalisations that is consistent within individuals across call types but variable across individuals. This last solution is well suited for species that continuously modify the vocalisations they produce. It should be noted that such a voice-print differs from a vocal signature in that it is likely not actively produced, but is a by-product of the vocal tract. To distinguish between these types of vocal signals, throughout this study we use the term ‘individual signature’ to denote actively produced uniqueness within call types and ‘voice-print’ to denote the emergent individual signature resulting from vocal tract morphology.

**Figure 1:**
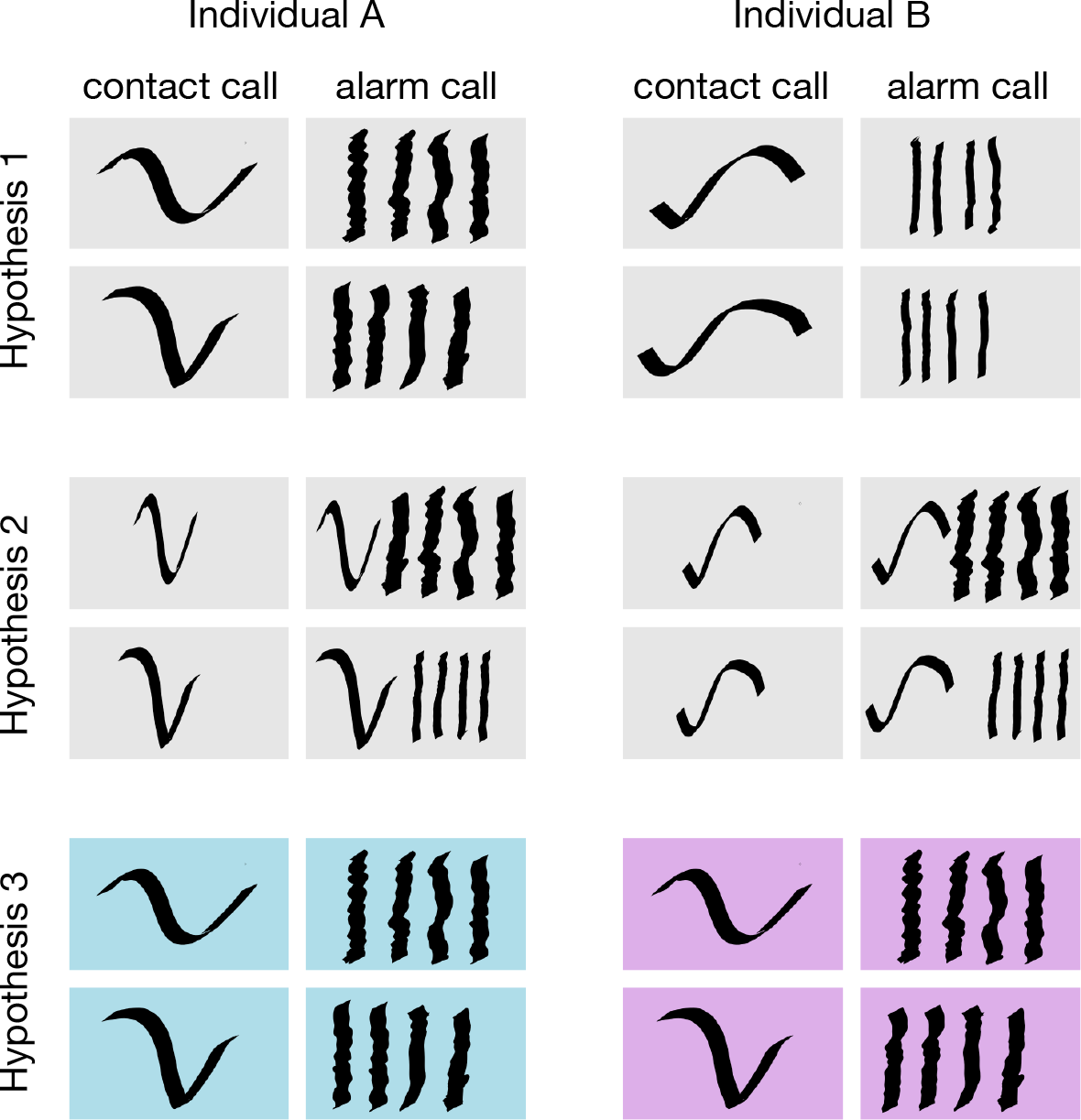
Illustration of how animals can encode individual signature in two call types. Squares are stylised spectrograms of contact and alarm calls. Rows within each hypothesis represent different versions of both call types for each individual. Hypothesis 1: Each call type is distinct -the individual ID is encoded in the frequency modulation of the contact call and the pulse duration of the alarm call. Hypothesis 2: An dedicated identity call -individual ID is only encoded in the frequency modulation of the contact call. The alarm call is now a sequence of contact-alarm to encode both individual ID and call function. The alarm call can be highly variable within individuals. Hypothesis 3: A voice-print -there is no individual information encoded in the frequency modulation or the pulse duration, but instead there is a general voice-print (represented by colour) that goes across call types.

The best known example of this third strategy is the voice-print in humans. Humans have a complex communication system with an almost limitless number of sounds that can be produced, rendering it unfeasible to include identity calls in combination with secondary utterances. Yet despite this flexible production, the human vocal tract leaves an individually distinct cue in the timbre of the voice, allowing speakers to be recognised across most utterances (Mathias and Kriegstein 2014). To date, the potential for such a voice-print to occur in other animals has received surprisingly little attention. Thus far, voice-prints have only been shown in the mating calls of red deer stags (*Cervus elaphus*), where Reby et al. (2006) used mel frequency cepstral coefficients combined with a hidden Markov chain model to find that 63% of roars and barks could be correctly assigned to seven individuals. Notably, this study used relatively few call types and individuals of a fixed repertoire species. To our knowledge there has been no study investigating voice-prints across call types in a non-human vocal learner with a large and flexible vocal repertoire. This is despite the fact that these species would benefit most from such an individual vocal recognition mechanism, since they might modify their contact call and thereby render an individual signature in frequency or duration less clear. Identifying if and what other species exhibit similar voice-prints is an important first step in understanding how vocal learning can evolve without obscuring individual identity information in the vocalisations.

Parrots are open-ended vocal production learners that often exhibit large and flexible vocal repertoires (Bradbury and Balsby 2016; Wright and Dahlin 2018). In this group, most research focus has been on contact calls, loud calls often made during group fusion events, or when individuals are isolated. These contact call are likely socially learned in early stages of development (Berg, Delgado, Cortopassi, et al. 2012; Teixeira et al. 2021) and are generally assumed to broadcast identity (Wright 1996; Smith-Vidaurre, Araya-Salas, and Wright 2020). Some species also appear to actively modify their contact call over periods of weeks to converge with pairs or with flock mates (Dahlin et al. 2014; Scarl and Bradbury 2009), and there is even evidence for rapid convergence within vocal exchanges (Wright, Hara, et al. 2015; Balsby and Bradbury 2009; Balsby, Momberg, and Dabelsteen 2012). Despite this flexibility, some species have a stable individual signature in their contact call, at least within the time period of focus (Thomsen, Balsby, and Dabelsteen 2013; Smith-Vidaurre, Araya-Salas, and Wright 2020; Berg, Delgado, Okawa, et al. 2011). Additionally, other species have a stable group level signature in their contact call that also appears to persist over long periods of time. For example yellow-naped amazons (*Amazona auropalliata*) have dialects that are virtually unchanged throughout a period of 11 years in some locations (Wright, Dahlin, and Salinas-Melgoza 2008). However, it is not known how much of an individual signature exists in call types other than the contact call for adult parrots (but see Wein et al. (2021)), whether this is stable over time, or if vocal distinctiveness carries across call types as a voice-print.

In our study we addressed these questions in monk parakeets (*Myiopsitta monachus*), a communal nesting parrot with a large native and invasive range. Monk parakeets are popular pets with good vocal imitative abilities and like all parrots, are life-long vocal learners. Their contact calls have been extensively studied (Martella and Bucher 1990; Buhrman-Deever, Rappaport, and Bradbury 2007; Smith-Vidaurre, Araya-Salas, and Wright 2020; Smith-Vidaurre, Perez-Marrufo, and Wright 2021; Smeele, Tyndel, et al. 2022), with these studies suggesting that monk parakeet contact calls contain an individual signature (Smith-Vidaurre, Araya-Salas, and Wright 2020). In their invasive range, they also appear to exhibit geographically distinct dialects in contact calls (Buhrman-Deever, Rappaport, and Bradbury 2007; Smeele, Tyndel, et al. 2022), although this is much less pronounced in their native range (Smith-Vidaurre, Araya-Salas, and Wright 2020). However, it should be noted that no study has recorded vocalisations from a large set of individually-marked monk parakeets, or extended this analysis to other call types. Here, we recorded 229 wild, individually-marked monk parakeet in Barcelona, Spain over a period of two months across two consecutive years, and manually categorised calls into 11 call types. First, for the five call types with enough data, we measured similarity between calls within the same call type and analysed the results with a Bayesian multilevel model to test how much individual signature exists in the most common monk parakeet call types and how stable these signatures are over time. Second, we tested how much individual information exists across call types by training the model on one set of call types and predicting on another set of call types. Based on previous work we predicted high levels of individual signature in contact calls and lower levels in other call types. Additionally, we predicted a stable signature over a month long period with reduction in similarity over years. Finally, if monk parakeets exhibit a voice-print in their vocalisations, we predicted that calls could be assigned to individuals across call types.

## Methods

### Study System

We studied monk parakeets in Parc de la Ciutadella and surrounding areas in Barcelona, Spain, where they have been reported as an invasive species since the late 1970s (Batllori and Nos 1985). Parc de la Ciutadella, Promenade Passeig de Lluís Companys and Zoo de Barcelona form a continuous habitat of grass and asphalt with multiple tree species in which monk parakeets nest and forage. They build complex stick nests in trees and other structures, often building new nest chambers on top of already existing nest structures (Eberhard 1998), creating colonies of birds living in close proximity.

Since May 2002, adults and juveniles have been regularly captured and marked using a walk-in trap on Museu de Ciències Naturals de Barcelona, while fledglings have been marked directly at their nests (Senar, Carrillo-Ortiz, and Arroyo 2012). Birds are ringed with unique leg-bands and fitted with neck collars with small tags displaying unique combinations of letters and digits. These are similar to small dog tags and can be read from up to 30 meters with binoculars. This effort has resulted in over 3,000 ringed birds since May 2002, of which 300-400 are recaptured/sighted each year. In November 2021, to increase the number of marked birds in the population for this study, we captured and tagged an additional 59 adults and juveniles at their nests, trapping individuals at night with hand nets. All birds were ringed with special permission EPI 7/2015 (01529/1498/2015) from Direcció General del Medi Natural i Biodiversitat, Generalitat de Catalunya, and with authorization to JCS for animal handling for research purposes from Servei de Protecció de la Fauna, Flora i Animal de Companyia (001501-0402.2009).

### Data Collection

Vocalisations were recorded from marked individuals in two years between 27.10.20 -19.11.20 and 31.10.21 -30.11.21 (55 days total) using a Sennheiser K6/ME67 shotgun microphone and Sony PCM D100 recorder from a distance ranging between one and 20 meters. The IDs and behaviours of focal individuals, the behaviours of close-by individuals and the general contexts of the vocalisations were verbally annotated. Some recordings were also videotaped and IDs were transcribed afterwards.

In addition, we mapped all nests in the recording area using Gaia GPS on several Android cellphones. Errors were manually corrected to less than 10 meters. In order to determine nest occupancy, we monitored nests multiple times throughout the day until an individual was observed inside the nest at least three times. Individuals were assigned to a nest entry if they were seen at least once inside one of the nest entrances. If they were sighted at multiple nests, they were assigned to the nest where they were most often sighted. If no birds were observed at a nest, we continued to monitor the nest daily for the duration of the recording period.

## Data Processing

All calls with fundamental frequencies clearly distinguishable from background noise and with no overlapping sounds were selected in Raven Lite (Cornell Lab of Ornithology, NY 2016). Calls were then manually assigned to 11 broad call types based on structural similarity. For five of these we had a large enough sample size to analyse the individual signature. These were: (1) contact call -a frequency modulated call with at least three infliction points, (2) *tja* call -a tonal call with a single rising frequency modulation, (3) *trruup* call -a combination of amplitude modulated introduction (similar to alarm calls) with a tonal ending (similar to the *tja* call), (4) alarm -an amplitude modulate call with at least four ‘notes’ and clear harmonics, predominantly used in distress situations, and (5) growl -an amplitude modulate call with at least four ‘notes’ and no clear harmonics, predominantly used in social interactions (see Figure 2). Other call types were included for the cross call type analysis (see further down).

**Figure 2:**
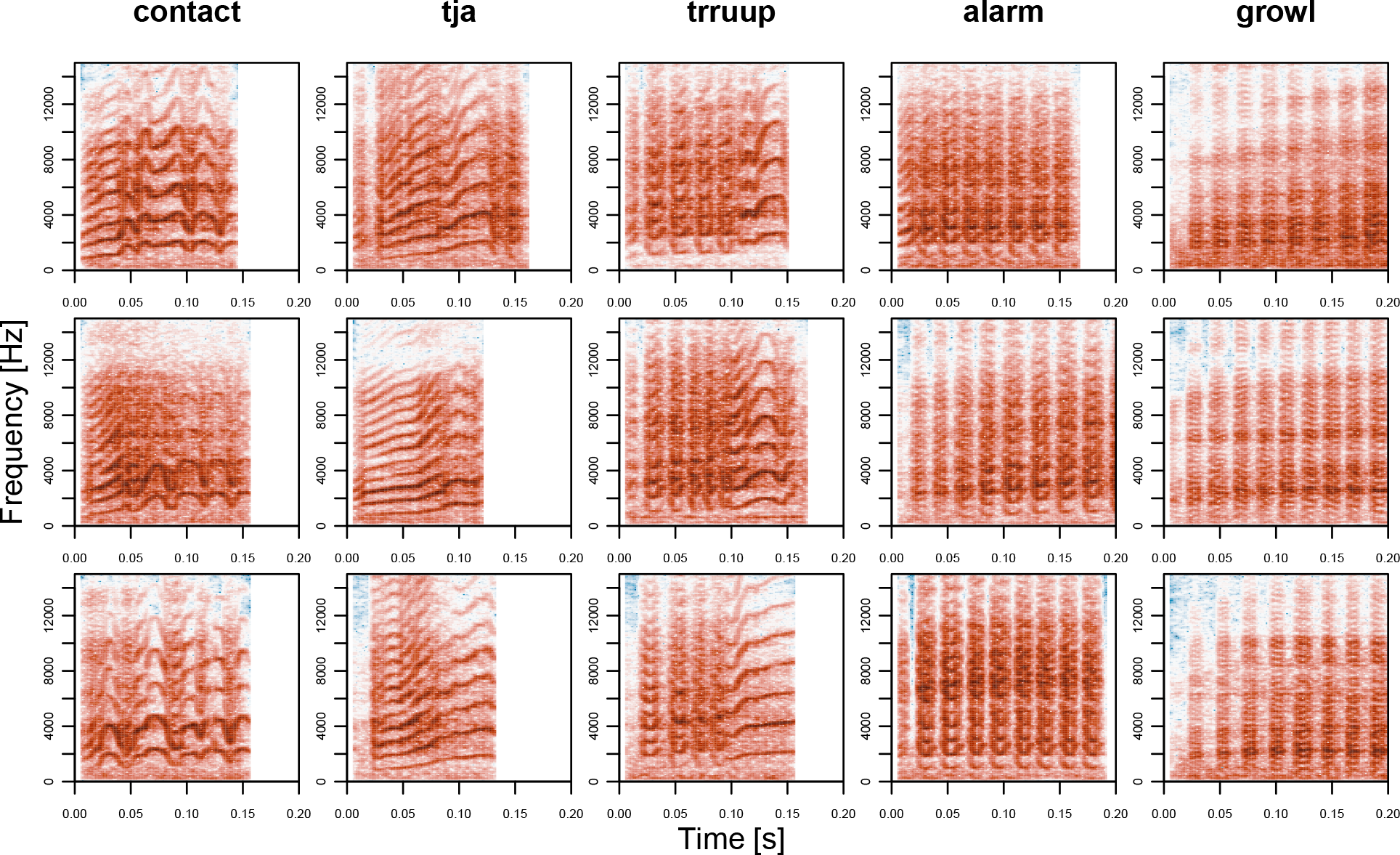
Example spectrograms of the call types included in the analysis of vocal signature. Settings: window length = 512, overlap = 89%, window type = Hanning. Darker colours (red) indicate more energy for that frequency (y-axis) at that time (x-axis).

We used four methods to measure similarity between calls: dynamic time-warping (DTW, (Giorgino 2009)), spectro-graphic cross correlation (SPCC, (Clark, Marler, and Beeman 1987)), spectrographic analysis (SPECAN, specified in the supplemental materials) and mel frequency cepstral coeffienct cross correlation (MF4C, specified in the supplemental materials). We present the results of SPCC in the main text, since SPCC could be run on all call types, is the most used method in previous work and other methods gave similar results. The results of all other methods are presented in the supplementary materials. SPCC consists of sliding two spectrograms over each other and calculating the sum of the difference between each pixel per sliding window. The distance at maximal overlap between calls is then used as a measure of acoustic distance (see Figure 3A for a schematic overview). We implemented our own function for SPCC in R (R Core Team 2021) to remove as much background noise as possible (see supplemental materials for details).

**Figure 3:**
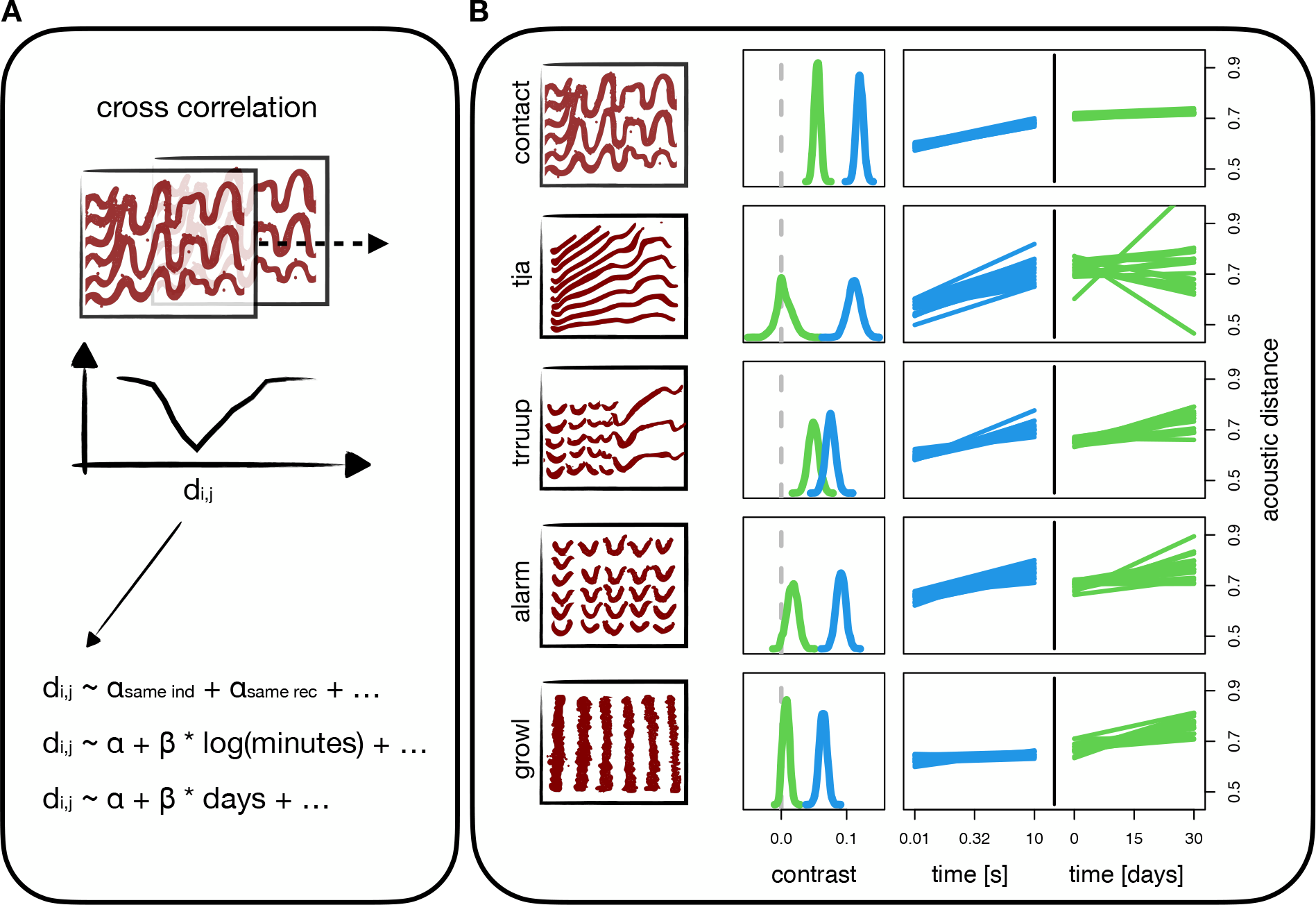
Workflow and results for spectrographic cross correlation. Black squares with thick red lines are stylised version of the spectrograms. For real data see Figure 2. (A) Schematic overview of the analysis pipeline. (B) Model results for contact calls, tja calls, trruup calls, alarm calls and growls (top to bottom). Blue density plots are the posterior contrast between the similarity of calls from different individuals versus the same individual and same recording. Green density plots are the posterior contrast between the similarity of calls from different individuals vs the same individual but different recordings. Little or no overlap with zero (dashed grey line) indicates a reliable signature of individual or recording. Blue lines are 16 samples from the posterior prediction of acoustic distance throughout time within a recording. Green lines are 16 samples from the posterior prediction of acoustic distance throughout days between a recording.

### Statistical Analysis

The first aim of this study was to determine whether call types contained an individual signature. Three of our methods (DTW, SPCC and MF4C) produce similarity matrices rather than single or multiple measures per call. The analysis of such a matrix is challenging, since most conventional models are designed for multivariate data sets. To estimate similarity between calls coming from the same individual compared to calls coming from different individuals, we therefore used a Bayesian model that is structurally similar to the social relationships model (Kenny and La Voie 1984). The response variable were dyadic acoustic distances and predictor variables were whether or not the calls came from the same individual, from the same recording, a unique ID for the recording dyad, a unique ID for the individual dyad, and a unique ID for both calls. This way we controlled for repeated and unbalanced sampling per individual, per recording and repeated comparisons per call (see supplemental materials for the mathematical model definition).

To visualise similarity between calls coming from the same versus different individuals, we computed the posterior contrast between the predicted acoustic distance between calls from two different individuals and between calls from two different recordings of the same individual. A contrast is the pairwise difference between samples of two distributions. This creates a new posterior distribution that reflects the modelled difference between two categories, in this case same vs different individual. We report the whole posterior density and the fraction of posterior samples that overlap with zero. If the contrast does not overlap zero, or there is only little overlap, it indicates that, given the data and model structure, there is a difference between categories. To visualise similarity between calls from the same recording session, we computed the posterior contrast between calls from two different individuals and compared that to posterior contrasts between calls from the same individual and same recording.

The second aim was to test how stable the individual signatures were across time. We tested this across three scales: within a recording, across days and across years. We only used acoustic distances between calls from the same individual. We then modeled the acoustic distance as a function of time separating the two calls with a Bayesian multilevel model (see supplemental materials for the full model definition). For the first model we included time on the log-scale. For the latter two models we only included acoustic distance between calls coming from different recordings and time was measured as days between recordings and same or different year respectively (see supplemental materials for the mathematical model definition).

Third, to assess how recognisable individuals were across call types we ran multiple permuted discriminant function analyses (pDFAs) on the mel frequency cepstral coefficients (MFCC) summary statistics (mean and standard deviation). We chose to write our own function to run pDFA in R (R Core Team 2021), so we could choose vocalisations from different recordings for the training and test sets, balance these data sets and compare the resulting scores to scores from a randomised data set. This function was based on the work done by (Mundry and Sommer 2007). To test how reliable pDFAs could score individual identity within a call type, we first trained and tested a pDFA on contact calls. To test how much information was available across broad call type categories we trained a pDFA on amplitude modulated calls (with clear interruptions in the amplitude -see Figure 2 alarm, *trruup* and growl for examples) and then tested on tonal calls (with uninterrupted tonal components, see Figure 2 contact and *tja* for examples) and vice versa (see Figure 4 for a schematic overview). We grouped call types to obtain a large enough sample size for the pDFA and choose these categories to maximise dissimilarity between the two categories. For all pDFAs we report the 89% highest density interval of differences between the trained and randomised score. We also report the overlap with zero. If there is no overlap zero, or the overlap with zero is very limited, it means the trained pDFA was performing above chance-level. To test if the model learned features related to sex or background noise we reran the procedure on calls from a females from Promenade Passeig de Lluís Companys, which is generally more noisy and also reran the procedure where labels were restricted to be randomised within location (Promenade Passeig de Lluís Companys and Parc de la Ciutadella). Throughout the text we use pDFA to refer to a full set of permuted discriminant function analyses and DFA to refer to a single run of discriminant function analyses.

**Figure 4:**
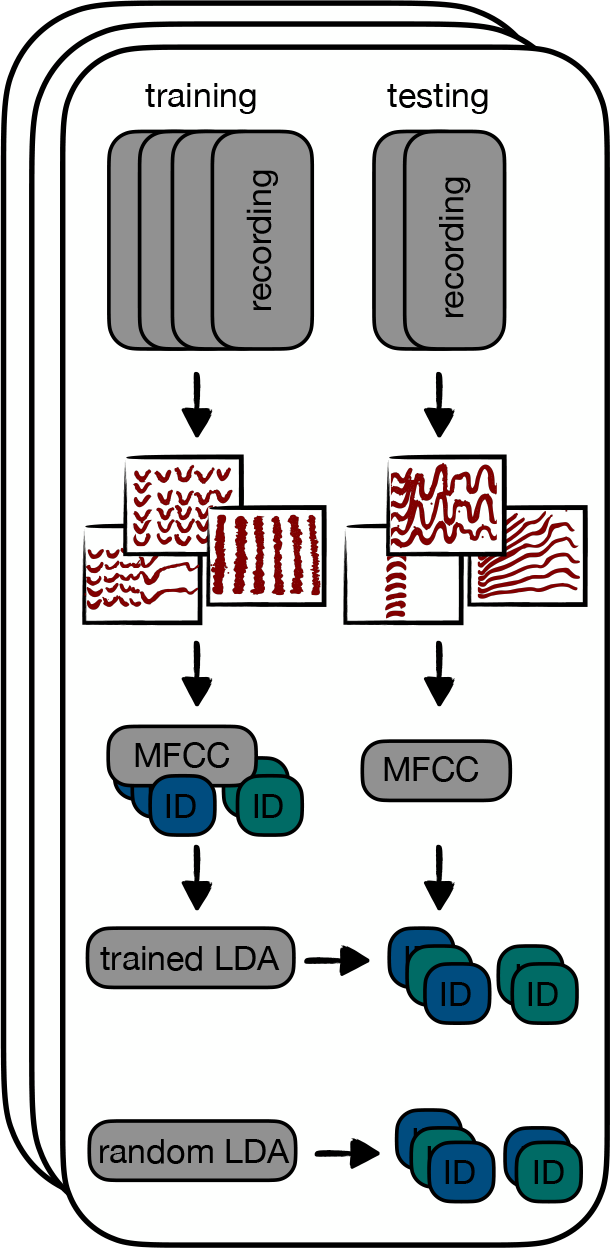
Workflow for permuted linear discriminant analysis (LDA) across call types. For each iteration recordings are split in a training and a testing set. A linear discriminant classifier is trained on mel frequency cepstral coefficients from amplitude modulated calls from the training set with calls labeled with individual ID. A random LDA, where labels are randomised is also trained. The correct classification percentage is calculated for the tonal calls from the testing set for both trained and random LDA. Black squares with thick red lines are stylised versions of the spectrograms.

All analysis was run in R (R Core Team 2021) and scripts are publicly available on GitHub: https://github.com/simeonqs/Evidence_for_vocal_signatures_and_voice-prints_in_a_wild_parrot. All Bayesian models were run using the R package *cmdstanr* (Gabry and Češnovar 2022), which runs the Stan sampler (Gelman, Lee, and Guo 2015). Rhat values were monitored to ensure convergence.

## Results

In total, we recorded 5599 calls across 229 individually marked birds over the two years of data collection, 3242 in year 1, and 2357 in year 2. Our manual sorting lead to 3203 contact calls, 185 *tja* calls, 265 *trruup* calls, 249 alarm calls and 364 growls. We then asked whether the five call types were individually distinctive. As expected from previous studies (Smith-Vidaurre, Araya-Salas, and Wright 2020), we found a weak but reliable individual signature for the contact call (contrast mean: 0.06, overlap zero: 0.00, Figure 3B). This contrast means that calls from the same individual are 0.06 closer to each other on the normalised scale (0 being completely similar, 1 being completely dissimilar) than calls from two different individuals. The *trruup* call contained an equally strong individual signature (contrast mean: 0.05, overlap zero: 0.00, Figure 3B). The individual signature in alarm calls was relatively weaker (contrast mean: 0.02, overlap zero: 0.02, Figure 3B). Finally, for the *tja* and growls there was no evidence for an individual signature (see Figure 3B).

Additionally, we found evidence in all call types for short-term temporal variability, with calls from the same recording sounding more similar than calls coming from two different recordings. For all calls other than the growl there was also an increase in acoustic distance with time throughout a recording (see Figure 3B). In other words, calls coming right after each other were more similar than calls spaced further apart in the recording. For the *trruup* call, alarm call and growl acoustic distance also increased with days between recordings. However, at the largest time scale this temporal variability disappeared, with individual signature stable between years and calls not more similar within year than across (see Figure 5). Our method did not allow us to test which spectral features (e.g., fundamental frequency or duration) changed most over time, since models were based on distance metrics (in other words comparing two whole calls to each other, rather than several metrics for each call).

**Figure 5:**
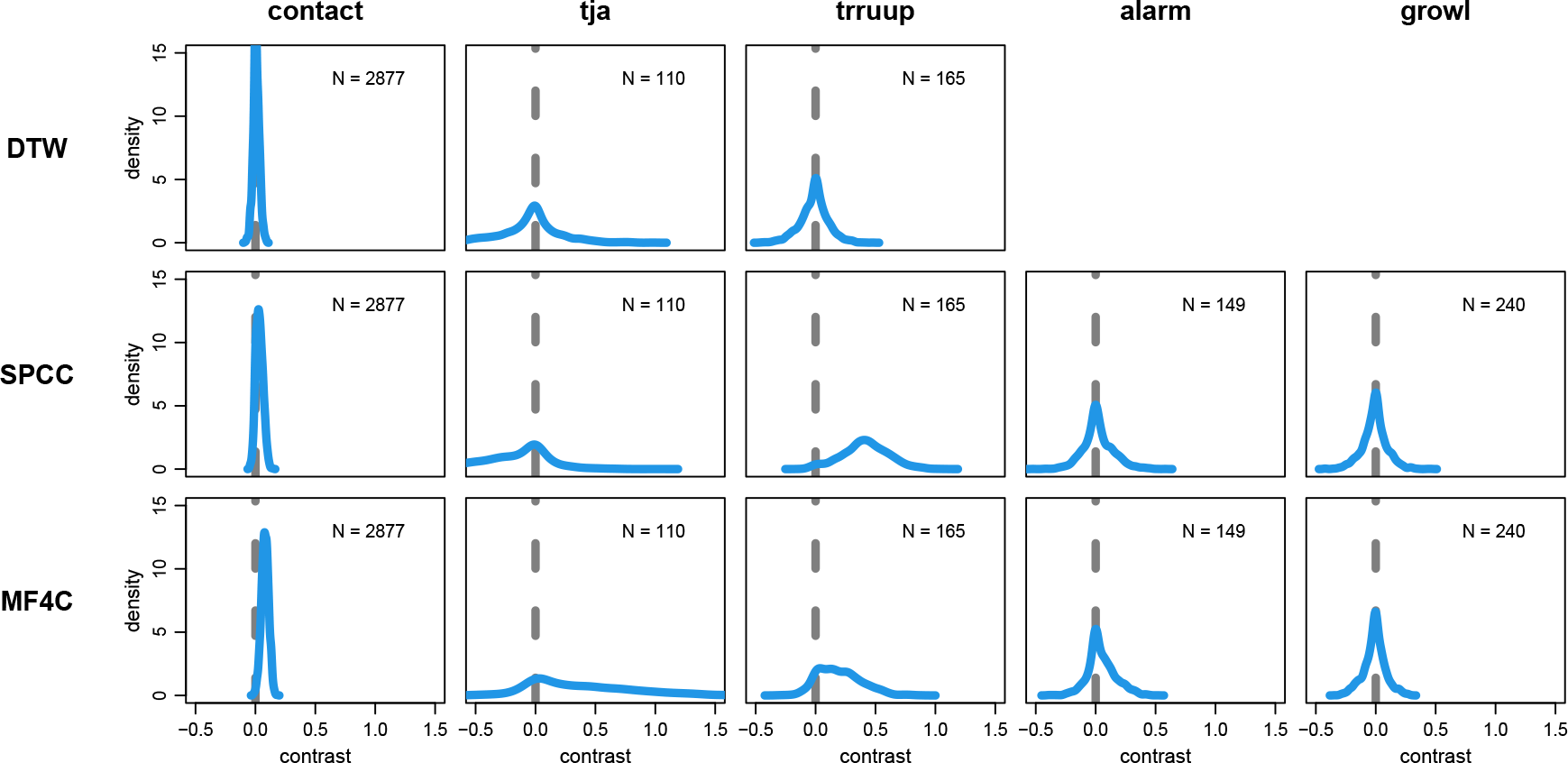
Posterior distributions for the for the Bayesian multilevel model with contrasts between recordings from the same vs different years. Greater values mean more within year similarity. Little to no overlap with zero (grey dashed line) indicate reliable differences. Models were run separately for the main call types (columns) and methods (rows; DTW -dynamic time warping, SPCC -spectrographic cross correlation, MF4C -mel frequency cepstral coefficient cross correlation). N represents the number of calls included in the model.

We then used multiple permuted discriminant function analyses (pDFAs) on the mel frequency cepstral coefficients summary statistics (mean and standard deviation) to test whether DFAs trained on a subset of calls were able to successfully predict caller identity when presented with new calls. First, and as expected, results from the pDFAs further added to the evidence that contact calls contained an individual signature, with the trained DFA was on average 35% more successful in predicting identity than a randomised DFA (see Table 1). We also found evidence that calls contain general individualised features that were maintained across call types. A pDFA with amplitude modulated calls as training data and tonal calls as testing data or vice versa achieved a score of 16% and 10% more successful respectively than random (see Table 1). The trained DFA outperformed the random DFA in all iterations of the model.

**Table 1:**
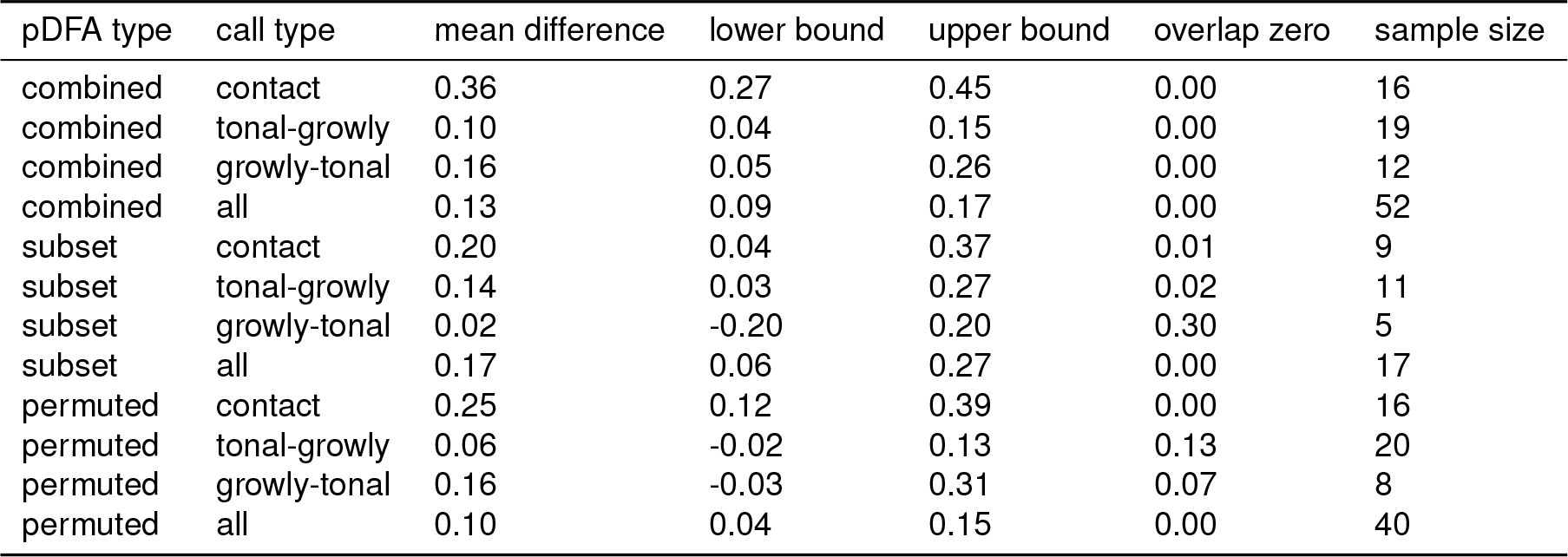
Table of all the results of permuted discriminant function analysis. The column ‘pDFA type’ contains information about how the pDFA was run: ‘combined’ included all recordings, ‘subset’ included only females in Promenade Passeig de Lluís Companys and ‘permuted’ was run where randomisation was done within location. The column ‘call type’ contains information about which call types were included. E.g., ‘tonal-growly’ means the model was trained on tonal calls and tested on amplitude modulated calls and ‘contact’ means it was trained and tested on contact calls. The column ‘mean diff’ contains the mean difference between the trained and random DFA’s. The column ‘low bound’ contains the lower bound of the 89% highest density interval. The column ‘up bound’ contains the upper bound of the 89% highest density interval. The column ‘overlap zero’ contains the fraction of iterations that were less than zero. The column ‘sample size’ contains the number of individuals included.

| pDFA type | call type | mean difference | lower bound | upper bound | overlap zero | sample size |
| --- | --- | --- | --- | --- | --- | --- |
| combined | contact | 0.36 | 0.27 | 0.45 | 0.00 | 16 |
| combined | tonal-growly | 0.10 | 0.04 | 0.15 | 0.00 | 19 |
| combined | growly-tonal | 0.16 | 0.05 | 0.26 | 0.00 | 12 |
| combined | all | 0.13 | 0.09 | 0.17 | 0.00 | 52 |
| subset | contact | 0.20 | 0.04 | 0.37 | 0.01 | 9 |
| subset | tonal-growly | 0.14 | 0.03 | 0.27 | 0.02 | 11 |
| subset | growly-tonal | 0.02 | -0.20 | 0.20 | 0.30 | 5 |
| subset | all | 0.17 | 0.06 | 0.27 | 0.00 | 17 |
| permuted | contact | 0.25 | 0.12 | 0.39 | 0.00 | 16 |
| permuted | tonal-growly | 0.06 | -0.02 | 0.13 | 0.13 | 20 |
| permuted | growly-tonal | 0.16 | -0.03 | 0.31 | 0.07 | 8 |
| permuted | all | 0.10 | 0.04 | 0.15 | 0.00 | 40 |

While we did our best to select calls with no overlapping features or background noise, it is possible that our analysis was still detecting features that were more likely to occur in calls of particular individuals. Alternatively, individuals might have called in a characteristic way in particular locations, creating a false signal in the data. To try and remove these potential biases, we re-ran our analysis within females in Promenade Passeig de Lluís Companys. In this case, only the pDFA trained on tonal calls and tested on amplitude modulated calls performed better than random (see Table 1). As this might be an effect of the greatly reduced dataset, we then re-ran our analysis with the full dataset, but restricting randomisation to only within location. In this case, the trained pDFA performed much better than chance, but overlap with zero increased to 13% and 7% for tonal to amplitude modulated and vice versa respectively (see Table 1).

## Discussion

Many animals are likely to benefit from individual recognition. In many species of birds, this is thought to most likely occur through individually distinct vocalisations. Yet how this is achieved in species with open-ended vocal production learning, and in parrots in particular, has been understudied (but see Thomsen, Balsby, and Dabelsteen (2013)). By recording vocalisations in individually-marked wild monk parakeets across one month and over two years, we reveal multiple insights into the vocal production of this parrot. First, we show that multiple call types given by monk parakeets contain a weak individual signature, but that this signature is relatively stable over time, persisting within and between years (see Figure 3 and supplemental materials). Second, we show that calls are not stereotyped, but are highly variable over short time scales (seconds to minutes, Figure 3B); within the same recording calls are generally more similar than calls from different recordings, and even within a recording calls close in time are more similar. Third, we tested if individual identity was distinguishable across call types. We used mel frequency cepstral coefficients (MFCC), training a permuted discriminant function (pDFA) on one set of call types and testing on another set of call types, doing so across recordings to make sure background noise could not be ‘learned’ by the model. Our results suggest monk parakeets have a voice-print that exists across structurally different call types, although the strength of evidence varied across call types and analyses. To our knowledge this is the first evidence for the detection of voice-prints in a non-human vocal learner.

The ability to recognise individuals from their vocalisations should be highly advantageous in species with social systems like monk parakeets, where individuals may encounter many potential association partners during fission-fusion foraging dynamics. Previous studies have demonstrated individual signatures in the contact calls of monk parakeets (Smith-Vidaurre, Araya-Salas, and Wright 2020), as well as in contact calls from other parrot species (Thomsen, Balsby, and Dabelsteen 2013; Balsby and Adams 2011; Farabaugh, Linzenbold, and Dooling 1994; Berg, Delgado, Okawa, et al. 2011). However, like many parrots, monk parakeets have a large and variable vocal repertoire, and individuals might benefit from individual recognition in multiple call types. For example, individual level variation might be important for alarm calls which are generally used when individuals are agitated by each other or by external threats (Blumstein and Munos 2005). The *trruup* call is also often given in situation where flocks fission (SQS personal observation), in which case it may be important to know which conspecifics are about to fly away. In support of this prediction, we found that three of five tested call types (contact, alarm and *trruup*) in monk parakeets contained some evidence for an individual signature. While we found no evidence for individual distinctiveness in the growl or *tja* calls, it might be these calls do not require individual signatures: the *tja* call is often used in combination with other calls, and the growl is often used in close-range social interactions where identity might have already been established. Alternatively, it could be that these calls cannot support individual signatures: the *tja* is too short to allow for many unique variants, and the growl has no tonal structure in which identity information could potentially be encoded. This is in line with results found for chimpanzees (*Pan troglodytes*), where the short range pant grunts contained less individual variation than other calls (Mitani, Gros-Louis, and Macedonia 1996).

We proposed three hypotheses for how a vocal recognition system could be achieved in monk parakeets (see Figure 1). First, individuals could utilize individual signatures in several call types, unique to each call. Second, individuals could utilize a single unique signal that is added to the vocal sequences of multiple call types. Third, each individual could have a set of vocal features that are shared across all their calls, i.e., a voice-print. While our results provide evidence for an individual signature present in some call types (supporting the first hypothesis), calls were also highly variable. Overall, our results that a model trained on one individual signatures in one call type could help predict individual identity in another call type best supports the third hypothesis, that monk parakeets possess a voice-print that exists across call types, with a shared set of structural features that make them individually recognisable. Overall, we found most support for this third hypothesis. The individual signatures in the contact call were reliable, but decayed rapidly. However, a weak individual signature remained even across years. The fact that we did find a voice-print even across structurally very different tonal and amplitude modulated calls strongly suggests that this could be the dominant mode of recognition. Leroux et al. (2021) put forward a method to detect voice-prints across sequences of calls, something that might improve individual recognition in follow up studies that also include such sequences.

It should be noted that we used mel frequency cepstral coeficients (MFCCs) and summarised these using the mean and standard deviation of each cepstral. There are two potential issues with this approach. First, the mel frequency range was initially designed to represent how humans perceive sound (Chakraborty, Talele, and Upadhya 2014) and it can be argued that this method is not designed to detect voice-print in non-human vocalisations. However, we believe that it is suitable here, because the orange-fronted conure (*Eupsittula canicularis*), a slightly smaller parrot, has been shown to have a comparable hearing sensitivity curve to humans (Wright, Cortopassi, et al. 2003) and monk parakeets have their fundamental frequency between 1 and 2 kHz, which is higher than the human voice, but still within the band where the mel frequency filters have an effect (Martella and Bucher 1990). While individuals are probably able to detect more detailed information compared with our summary statistics, this can only currently be disentangled experimentally. For instance, future work could use play-backs to establish if and how well monk parakeets and other parrot species are able to recognise individuals across call types. Under this paradigm, and similar to Isabelle Charrier, Mathevon, and Jouventin (2002), one could potentially modify calls during playback to determine which spectral and temporal features are needed for individual recognition. Second, MFCCs can be sensitive to background noise. To deal with this, we ran several models to test how robust our results were when permuting the DFAs within location. We found that although performance decreased, there was still a clear trend for the trained DFA to outperform the random DFA. The drop in performance is likely a result of reduced sample size, and further studies are therefore needed to validate these results with a larger sample size.

It is also important to note that we cannot exclude the possibility that each call type also contains an individual signature in addition to the potential voice-print. However if parrots can learn to recognise individuals based on a voice-print shared across calls, such a generalised mechanism relaxes the pressure to produce structural components in each call. This allows calls to include other signatures (e.g., group identity) and reduces memory burden on the receiver significantly. There is also a good reason to expect voice-prints to be present in parrots. Unlike song birds, that produce their vocalisation using two relatively independent syringeal sound sources, parrots have only one sound source and modulate their vocalisations using trachea, tongue and beak. This is very similar to how humans produce the sounds that make up words (Nottebohm 1976; Ohms et al. 2012; Larsen and Goller 2002; Beckers, Nelson, and Suthers 2004; Patterson and Pepperberg 1994; Warren, Patterson, and Pepperberg 1996; Bottoni, Masin, and Lenti-Boero 2009; Brittan-Powell et al. 1997). This modulation or filtering by the vocal tract allows for many more individual-specific features to arise and make a voice-print more recognisable. Finally, a distinct and recognisable voice-print could be a particularly useful strategy used to manage individual recognition for species like parrots that are open-ended vocal learners living in complex but cohesive social groups.

Indeed, along with these main results, we found a high degree of variability within calls, with calls spaced ten minutes apart much less similar than calls spaced a second apart. It is unlikely that variation in background noise played a role in producing this result, since dynamic time warping performed on manually cleaned fundamental frequency traces obtained similar results (see supplemental materials). A more plausible explanation is that individuals are not capable of reproducing exactly the same call after too much time has elapsed. It is also possible that monk parakeets modify their call based on the context, audience, or emotional state. For example, some variants might be used in a foraging context where a partner is present while others are given in isolation. Another third possibility is that monk parakeets actively modify their contact call to match other individuals in their group, similar to the rapid convergence found in orange-fronted conures (Balsby and Bradbury 2009). If this is the case, we would expect a sequence of calls to vary depending on whom an individual is directing their call towards and the size of the audience. This would also suggest that individuals in larger groups should exhibit more variable calls. Both of these scenarios remain to be studied in more depth. However the presence of voice-prints may help explain how individuals can have such variable calls. If individuals modify the tonal structure of their contact calls in call response interactions, the individual signature in those calls will degrade over seconds within a recording. The voice-print would, however, be much more stable, given it’s generated by the morphology of the vocal apparatus, and it would still provide the conspecific with reliable features to recognise the vocalising individual.

The fact that individuals are so variable in their calls raises a methodological problem for dialect studies on unmarked populations. When recording in the wild, individuals can generally only be monitored for short periods of time. For example, in our study it was rarely possible to record individuals for more than 3-5 minutes. In this short period individuals are likely to exhibit a consistent individual signature, but this signature was less consistent across recordings. A common technique to exclude repeated sampling of individuals across recordings is to look for highly similar calls and exclude these (Buhrman-Deever, Rappaport, and Bradbury 2007; Smith-Vidaurre, Araya-Salas, and Wright 2020). However, this assumes one can reliably estimate how similar a call needs to be in order to classify it as the same individual. We show that this can not be reliably estimated from short-term recordings. Moreover, we show that determining which calls come from the same individual in a large sample is not realistic given the amount of individual variability in contact calls. Instead, we suggest estimating the probability of recording the same individual multiple times and using a sensitivity analysis to test if the detected dialect signal is likely to be a true signal, or if it could have been caused by pseudoreplication (see e.g., Smeele, Tyndel, et al. (2022)).

## Conclusion and Outlook

Despite decades of research, the ability of parrots to identify each other based on vocalisations is still not well understood. Some species have clear group signatures and dialects (Wright 1996), while others appear to have much more pronounced individual signature in their contact calls (Buhrman-Deever, Rappaport, and Bradbury 2007; Thomsen, Balsby, and Dabelsteen 2013; Smith-Vidaurre, Araya-Salas, and Wright 2020; Berg, Delgado, Okawa, et al. 2011). This study provides the first evidence for an individual voice-print across multiple call types in parrots. Additionally, it demonstrates significant individual variability in the contact call over recordings, but with sustained stability over time. Finally, our findings suggest that the contact call is not unique in its ability to broadcast caller identity in parrots. Instead it appears that parrots may have potentially evolved the capacity for individual recognition across multiple calls types (Lavan et al. 2019). While our study provides evidence for detectable voice-prints in monk parakeets, further investigation is needed to establish whether parrots actively use voice-prints to recognize conspecifics. More generally, it would now be exciting to test if voice-prints are present in other species as well, and, if these voice prints are used for recognition, to further explore the dynamics driving the evolution of voice-prints, including whether their presence is predicted by life-long vocal learning or complex social interactions.

## Data accessibility

Code and small data files will be publicly available on GitHub: https://github.com/simeonqs/ Evidence for vocal signatures and voice-prints in a wild parrot. The full repository including large data files will be made publicly available on Edmond upon acceptance. A pre-print was deposited at BioRxiv (Smeele, Senar, et al. 2023).

## Ethics

All monk parakeets were ringed and blood samples taken with special permission EPI 7/2015 (01529/1498/ 2015) from Direcció General del Medi Natural i Biodiversitat, Generalitat de Catalunya, following Catalan regional ethical guidelines for the handling of birds. JCS received authorization (001501-0402.2009) for animal handling for research purposes from Servei de Protecció de la Fauna, Flora i Animal de Companyia, according to Decree 214/1997/30.07.

## Acknowledgements

We would like to thank Zoo Barcelona and Josep M. Alonso Farré for granting us access to the zoo grounds and showing us around. We would like to thank Andrés Manzanilla, Gustavo Alarcón-Nieto, Mireia Fuertes Clavero, Alba Ortega-Segalerva and JoséG. Carrillo-Ortiz for all their effort during fieldwork. Finally, we would like to thank Francesca S. E. Dawson Pell, Daniel Redhead, Jack W. Bradbury, Susannah Buhrman-Deever, Grace Smith-Vidaurre, Timothy F. Wright, Roger Mundry, Mirjam Knörnschild and Elizabeth Hobson for valuable advice during the early stages of this project.

