## supplemental materials for "Evidence for vocal signatures and voice-prints in a wild parrot"

**References** **18**

### 1 Supplemental Methods

#### 1.1 Observer reliability

To test how reliable calls were scored into the 11 call types, a random subset of 200 calls were scored by a second observer with prior experience to monk parakeet call type classification. The second observer had access to three examples per call type (not included in the random set) and could assign each of the random calls to only one call type. After sorting nine calls were excluded because they did not belong to any of the 11 call types (they were sorted into a 'other' category by the first observer). For the remaining classifications we plotted a confusion matrix and report the accuracy from the function *confusionMatrix* from the package *caret* (Kuhn and Max 2008). We repeated this analysis for the binary classification into tonal and amplitude modulated calls.

#### 1.2 Quantification of similarity

For tonal calls the fundamental frequency was traced in Luscinia (Lachlan 2007). For non-tonal calls only the start and end times were determined manually. Traces were then exported as CSV and imported into R (R Core Team 2021) for further analysis. Traces were smoothed using two steps. The first step filled gaps within a call by connecting the fundamental frequency at both sides of the gap with a straight line. To get rid of small measurement jitter, we used smooth.spline from *stats* with spar = 0.1 as a second step.

To measure the similarity between any two calls a variety of methods have been used. The most common are 1) dynamic time warping (DTW), which measures measures similarity between the fundamental frequencies of two calls while controlling for differences in speech rhythm (Bellman and Kalaba 1959); 2) spectrographic cross correlation (SPCC), which measures similarity between the spectrograms of two calls (Clark, Marler, and Beeman 1987); and 3) spectrograpic analysis (SPECAN) which measures a set of variables on the spectrogram. We added a fourth method not commonly used and cross correlated the mel frequency cepstral coefficients (MF4C).

##### 1.2.1 Spectrographic cross correlation (SPCC)

We used a set of custom functions to run the SPCC. We used the raw audio files and the start and end times determined in Luscinia. We used the function *ffilter* from the package *seewave* (Sueur, Aubin, and Simonis 2008) to apply a high pass filter from 500 Hz before analysis. Then we created spectrograms with window length = 512, overlap = 89% and window type = Hanning, which were modified to:

- be limited to frequencies between 1-3 kHz
- have standardised pixel values (by substracting the mean of all pixels and deviding by the standard deviation)
- have all values below 1.1 on the standardised scale to be set to 0 (to remove noise)
- have all values above 1.8 on the standardised scale to bet set to 1.8 (to soften loud components)
- then have all pixels sum to 1 (deviding each pixel by the sum of all the pixels in the spectrogram - to ensure that long spectrograms would weigh as much as short)

These modified spectrograms were then run through a function that slid them over each other in the time dimension and computed the difference for each step. The minimal difference (the difference at point of maximal overlap of the signal) was then used as measure of acoustic distance.

##### 1.2.2 Dynamic time warping (DTW)

We used the function *dtw* from the package *dtw* (Giorgino 2009) to run DTW on the fundamental frequency traces. We normalised and log transformed the resulting distance matrix.

##### 1.2.3 Spectrographic analysis (SPECAN)

We ran spectrographic analysis on the raw audio files. We used the start and end times determined in Luscinia. We used the function *ffilter* from the package *seewave* (Sueur, Aubin, and Simonis 2008) to apply a bandpass filter between 500 and 6,000 Hz before analysis. We then measured the following variables:

- duration (s)
- peak frequency (kHz): the loudest frequency component from the spectrum over the whole signal
- low frequency (kHz): the lowest frequency above 0.4 times the loudest component from the spectrum over the whole signal
- high frequency (kHz): the highest frequency above 0.4 times the loudest component from the spectrum over the whole signal
- bandwidth (kHz): the difference between highest and lowest frequency
- number of amplitude modulation peaks (n): the number of peaks in the spectral envelope
- inter peak interval (s): the median time between amplitude modulation peaks

###### 1.2.4 Mel frequency cepstral coefficient cross correlation (MF4C)

We first created MFCC objects. These consisted of matrices with MFCC traces as rows. We used the raw audio files and the start and end times determined in Luscinia. We used the function *ffilter* from the package *seewave* (Sueur, Aubin, and Simonis 2008) to apply a high pass filter from 500 Hz before analysis. We then used the *melfcc* function from the *tuneR* package (Ligges et al. 2018) to generate the traces. We used the following settings: wintime = 512/44100, hoptime = 50/44100, numcep = 10, minfreq = 300. We then used the same custom function as for SPCC to run the cross correlation.

##### 1.3 Statistical analysis

###### 1.3.1 Model I: individual and recording signal

We used the dis-similarity matrices from DTW, SPCC, MF4C or SPECAN to get a pairwise acoustic distance between all calls. We only included call pairs from the same year. We used a model similar to the Bayesian social relations model used in psychology. In our case the acoustic distance between any two calls was the response variable and was standardised before analysis. We included a global intercept and off-sets for: same vs different individual, same vs different recording, pair of individuals from which the calls came, pair of recordings from which the calls came and each call. The full model structure was as follows:

$$
\begin{aligned}
\text{accoustic distance} &\sim \text{normal}(\mu,\ \sigma) \\
\mu_n &= \bar{\alpha} + \alpha_{\text{same ind[n]}} + \alpha_{\text{same rec[n]}} + \\
&\quad \alpha_{\text{ind pair[n]}} + \alpha_{\text{rec pair[n]}} + \alpha_{\text{call i[n]}} + \alpha_{\text{call j[n]}} \\
\bar{\alpha} &\sim \text{normal}(0,\ 0.25) \\
\alpha_{\text{same ind}} &\sim \text{normal}(0,\ \sigma_{\text{same ind}}) \\
\alpha_{\text{same rec}} &\sim \text{normal}(0,\ \sigma_{\text{same rec}}) \\
\alpha_{\text{ind pair}} &\sim \text{normal}(0,\ \sigma_{\text{ind pair}}) \\
\alpha_{\text{rec pair}} &\sim \text{normal}(0,\ \sigma_{\text{rec pair}}) \\
\alpha_{\text{call}} &\sim \text{normal}(0,\ \sigma_{\text{call}}) \\
\sigma_{\text{same ind}},\ \sigma_{\text{same rec}},\ \sigma_{\text{ind pair}},\ \sigma_{\text{rec pair}} &\sim \text{exponential}(3) \\
\sigma,\ \sigma_{\text{call}} &\sim \text{exponential}(5)
\end{aligned}
$$

We fitted an un-centred version of the model using the package *cmdstanr* (Gabry and Češnovar 2021) with the No U-turn Sampler in Stan (Gelman, Lee, and Guo 2015) from R (R Core Team 2021). We ran 2000 iterations on four chains with adapt_delta = 0.99 and max_treedepth = 15. We used the difference between $\alpha_{\text{same ind [same]}}$ and $\alpha_{\text{same ind [different]}}$ as measure of individual signal and the difference between $\alpha_{\text{same rec [same]}}$ and $\alpha_{\text{same rec [different]}}$ as measure of recording signal. The other off-sets were included to control for unbalanced data and non-independence (e.g., some individuals might sound very much alike and we included multiple pairs of calls from the same two individuals).

###### 1.3.2 Model II: decay over time within recording

To test how stable the individual signature is within a recording we subsetted the pairwise dis-similarities to only include comparisons between calls from the same individual and recording. We included log10 of the time (in minutes) between the calls. We then used a multi-level model with a global intercept, a slope for the time-effect, off-sets for the intercept pair of individuals from which the calls came, pair of recordings from which the calls came and each call. We also included off-sets on the slope for individual. The full model structure was as follows:

$$
\begin{aligned}
\text{accoustic distance} &\sim \text{normal}(\mu,\ \sigma) \\
\mu_n &= \bar{\alpha} + \alpha_{\text{ind[n]}} + \alpha_{\text{rec[n]}} + \alpha_{\text{call i[n]}} + \alpha_{\text{call j[n]}} \\
&\quad (\bar{\beta} + \beta_{\text{ind[n]}} + \beta_{\text{rec[n]}}) * \text{time}_n \\
\bar{\alpha} &\sim \text{normal}(0,\ 0.5) \\
\alpha_{\text{ind}} &\sim \text{normal}(0,\ \sigma_{\text{ind}}) \\
\alpha_{\text{rec}} &\sim \text{normal}(0,\ \sigma_{\text{rec}}) \\
\alpha_{\text{call}} &\sim \text{normal}(0,\ \sigma_{\text{call}}) \\
\bar{\beta} &\sim \text{normal}(0,\ 0.3) \\
\beta_{\text{ind}} &\sim \text{normal}(0,\ \xi_{\text{ind}}) \\
\beta_{\text{rec}} &\sim \text{normal}(0,\ \xi_{\text{rec}}) \\
\sigma,\ \sigma_{\text{ind}},\ \sigma_{\text{rec}},\ \xi_{\text{ind}},\ \xi_{\text{rec}} &\sim \text{exponential}(2) \\
\sigma_{\text{call}} &\sim \text{exponential}(3)
\end{aligned}
$$

The model was fitted in the same way as model I.

###### 1.3.3 Model III: decay over days

To test how stable the individual signature is across days we subsetted the pairwise dis-similarities to only include comparisons between calls from the same individual and year, but from different recordings. We also included the difference in days between the calls. We then used a multi-level model with a global intercept, a slope for the time-effect, off-sets for the intercept for individual pair, recording and call, and off-sets on the slope for individual. The full model structure was as follows:

$$
\begin{aligned}
\text{accoustic distance} &\sim \text{normal}(\mu,\ \sigma) \\
\mu_n &= \bar{\alpha} + \alpha_{\text{ind pair[n]}} + \alpha_{\text{rec pair[n]}} + \alpha_{\text{call i[n]}} + \alpha_{\text{call j[n]}} \\
&\quad (\bar{\beta} + \beta_{\text{ind[n]}}) * \text{time}_n \\
\bar{\alpha} &\sim \text{normal}(0,\ 0.5) \\
\alpha_{\text{ind pair}} &\sim \text{normal}(0,\ \sigma_{\text{ind pair}}) \\
\alpha_{\text{rec pair}} &\sim \text{normal}(0,\ \sigma_{\text{rec pair}}) \\
\alpha_{\text{call}} &\sim \text{normal}(0,\ \sigma_{\text{call}}) \\
\bar{\beta} &\sim \text{normal}(0,\ 0.5) \\
\beta_{\text{ind}} &\sim \text{normal}(0,\ \xi_{\text{ind}}) \\
\sigma,\ \sigma_{\text{ind pair}},\ \sigma_{\text{rec pair}},\ \sigma_{\text{call}},\ \xi_{\text{ind}} &\sim \text{exponential}(2)
\end{aligned}
$$

The model was fitted in the same way as model I.

###### 1.3.4 Model IV: difference between years

To test how stable the individual signature is across years we subsetted the pairwise dis-similarities to only include comparisons between calls from the same individual, but from different recordings. We also included whether or not the recordings came from the same year. We then used a multi-level model with a global intercept, off-sets for: same vs different year, the individual, recording pair and call. The full model structure was as follows:

$$
\begin{aligned}
\text{accoustic distance} &\sim \text{normal}(\mu,\ \sigma) \\
\mu_n &= \bar{\alpha} + \alpha_{\text{same year[n]}} + \alpha_{\text{ind[n]}} + \\
&\qquad \alpha_{\text{rec pair[n]}} + \alpha_{\text{call i[n]}} + \alpha_{\text{call j[n]}} \\
\bar{\alpha} &\sim \text{normal}(0,\ 0.25) \\
\alpha_{\text{same year}} &\sim \text{normal}(0,\ \sigma_{\text{same year}}) \\
\alpha_{\text{ind}} &\sim \text{normal}(0,\ \sigma_{\text{ind}}) \\
\alpha_{\text{rec pair}} &\sim \text{normal}(0,\ \sigma_{\text{rec pair}}) \\
\alpha_{\text{call}} &\sim \text{normal}(0,\ \sigma_{\text{call}}) \\
\sigma,\ \sigma_{\text{same year}},\ \sigma_{\text{ind}},\ \sigma_{\text{rec pair}},\ \sigma_{\text{call}} &\sim \text{exponential}(3)
\end{aligned}
$$

The model was fitted in the same way as model I.

###### 1.3.5 pDFA

To test if there are features that contain an individual signal across call types we used mel frequency cepstral coefficients (MFCC). More specifically we used the function *melfcc* from the package *warbleR* (Araya-Salas and Smith-Vidaurre 2017) with the settings wintime = 512/44100, hoptime = 50/44100 and minfreq = 300 to retrieve the first ten cepstral coefficient traces. For each trace we saved the mean and standard deviation. These 20 measures per call were then used for the pDFA.

We wrote a custom R script to run the pDFA. We ran 100 iterations with the following steps:

- for each individual we randomly selected recordings until the set of recordings contained enough calls for testing (N_test)
- we then selected the other recordings for training
- if there were not enough recordings to fullfill both N_test and N_train, we did not include this individual
- for the individuals with enough data, we randomly selected N_test calls from the testing recordings and N_train calls from the training recordings
- we then used the standardised MFCC values to train an LDA (using the function *lda* from the package *MASS* (Venables and Ripley 2002))
- this LDA was then used to predict the IDs of the test data, the resulting proportion of correct classification was the trained score
- we repeated the previous two steps with both training and testing IDs randomised within the set and retrieved the random score
- for each iteration the difference between the trained and random score was reported

#### 2 Supplemental Results

##### 2.1 Observer reliability

The accuracy for the full call type classification was 0.72 (95% CI: 0.65, 0.78). Especially the short tonal calls and the soft growl were mis-classified (see Figure S1.

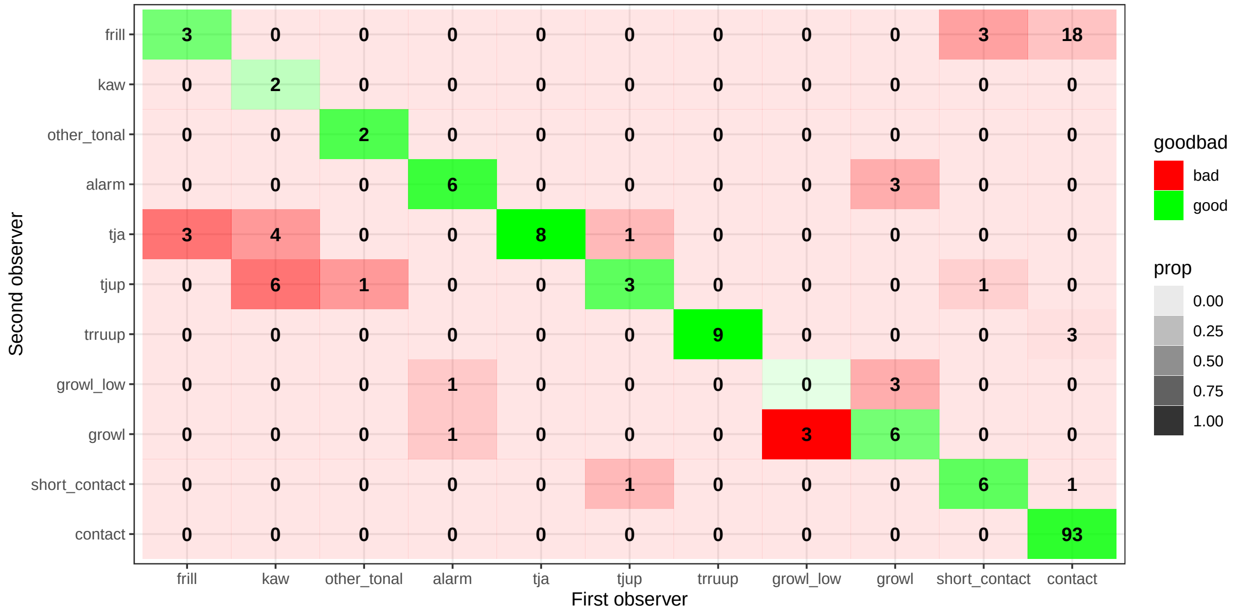

Figure S1: Confusion matrix for observer reliability. Numbers indicate how many calls were classified in that category, colour represent relative proportions.

The accuracy for the binary classification was 0.98 (95% CI: 0.95, 1.00). Only three growly calls were mis-classified (see Figure S2.

##### 2.2 Model I: boxplot raw data

To give an idea of the difference in similarity scores between calls coming from the same individual (but different recordings) and from different individuals we plotted boxplots for each of the five main call types (see Figure S3).

##### 2.3 Model I-III: results other methods

Results for dynamic time warping (see Figure S4) are similar to those of spectrographic cross correlation as presented in the main text. Note dynamic time warping cold only be run for tonal calls.

Individual signal was much reduced for spectrographic analysis (see Figure S5) compared to those of spectrographic cross correlation as presented in the main text.

Individual signal was much reduced for mel frequency cepstral coefficient cross correlation (see Figure S6) compared to those of spectrographic cross correlation as presented in the main text.

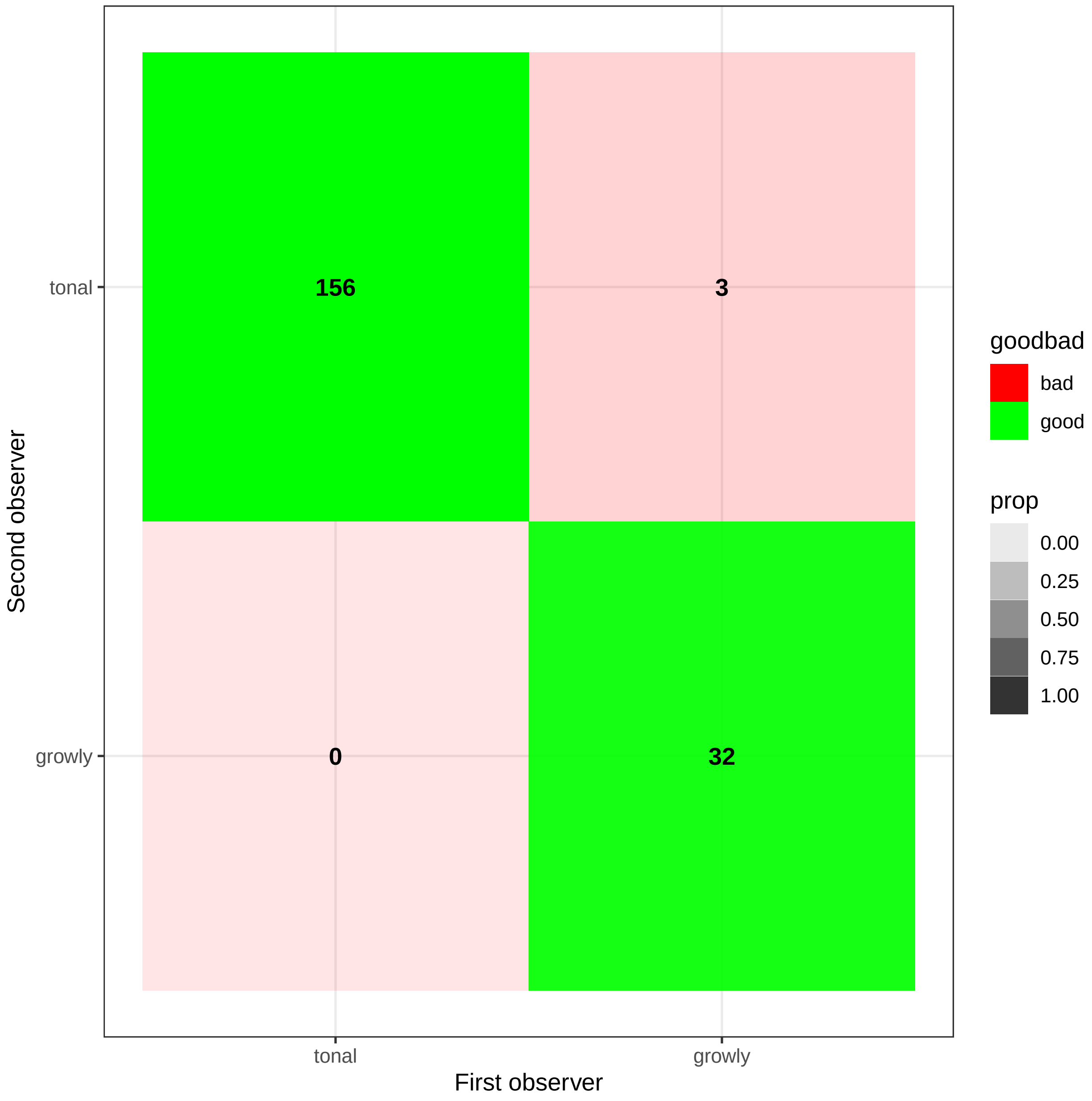

Figure S2: Confusion matrix for observer reliability with binary categories. Numbers indicate how many calls were classified in that category, colour represent relative proportions.

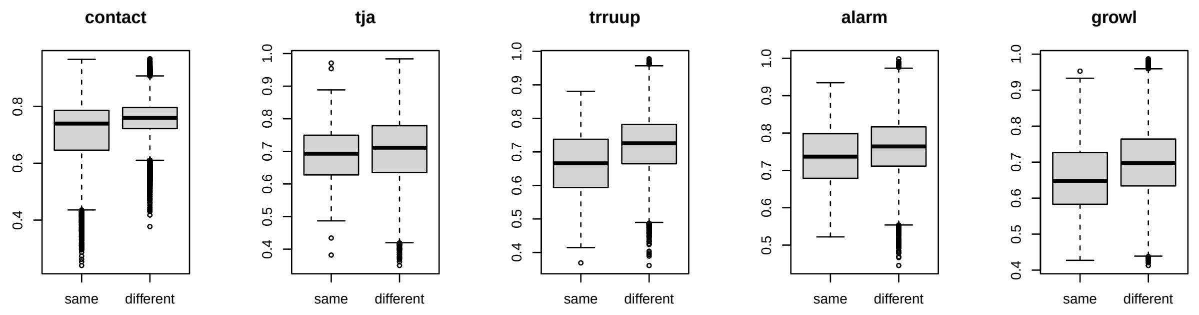

Figure S3: Boxplots for similarity (SPCC) between calls from the same or different individuals across the five main call types.

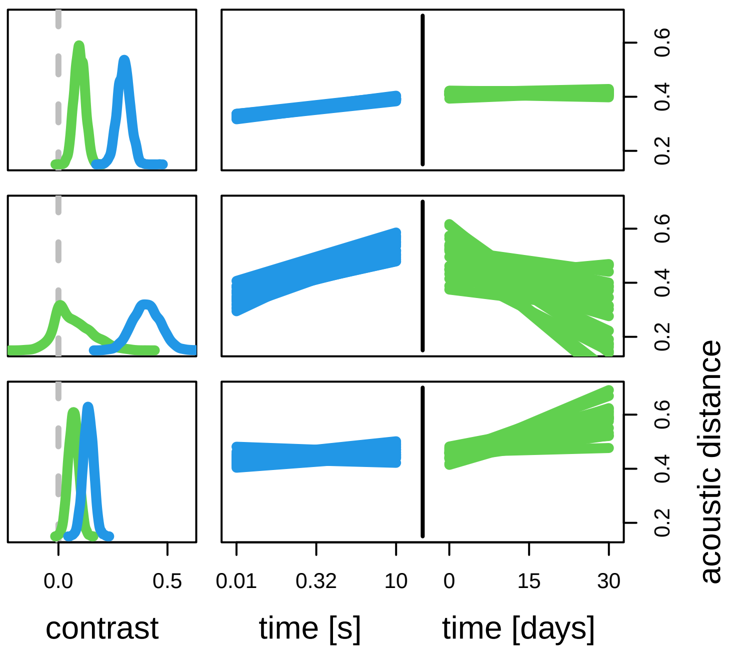

Figure S4: Model results for contact calls, tja calls amd trruup calls (ttb) using dynamic time warping. Blue density plots are the posterior contrast between the similarity of calls from different individuals versus the same individual and same recording. Green density plots are the posterior contrast between the similarity of calls from different individuals vs from the same individual but different recordings. Blue lines are 20 samples from the posterior prediction of acoustic distance throughout time within a recording. Green lines are 20 samples from the posterior prediction of acoustic distance throughout days between a recording.

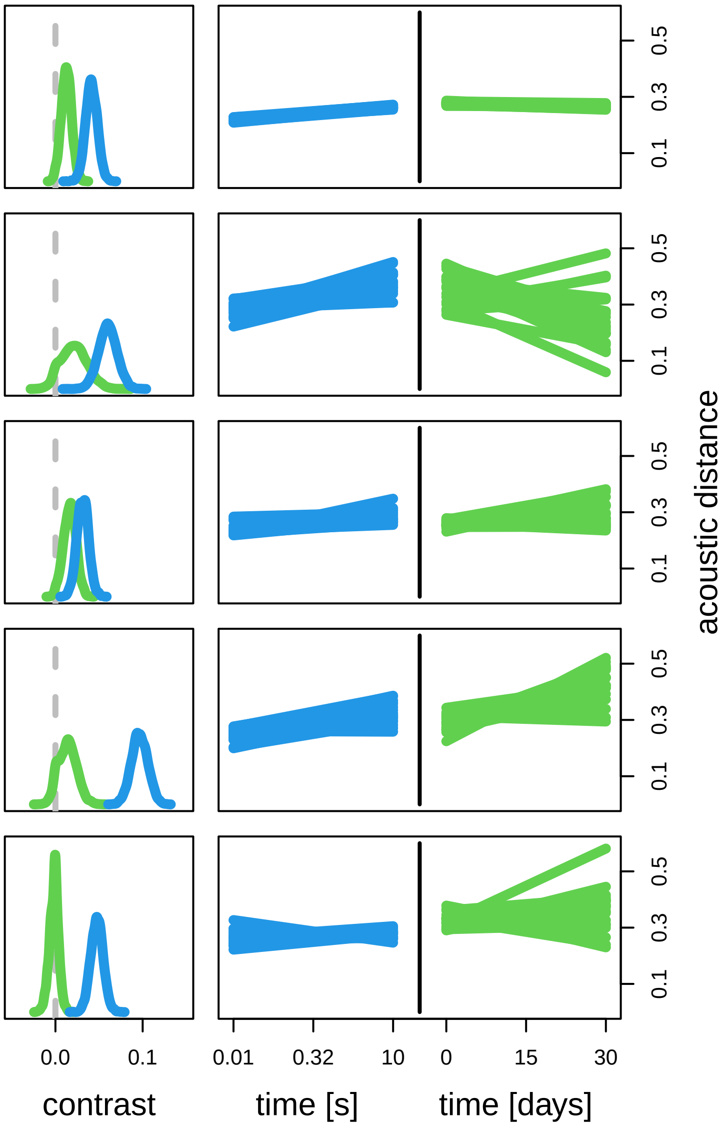

Figure S5: Model results for contact calls, tja calls, trruup calls, alarm calls and growls (ttb) using spectrogaphic analysis. Blue density plots are the posterior contrast between the similarity of calls from different individuals versus the same individual and same recording. Green density plots are the posterior contrast between the similarity of calls from different individuals vs from the same individual but different recordings. Blue lines are 20 samples from the posterior prediction of acoustic distance throughout time within a recording. Green lines are 20 samples from the posterior prediction of acoustic distance throughout days between a recording.

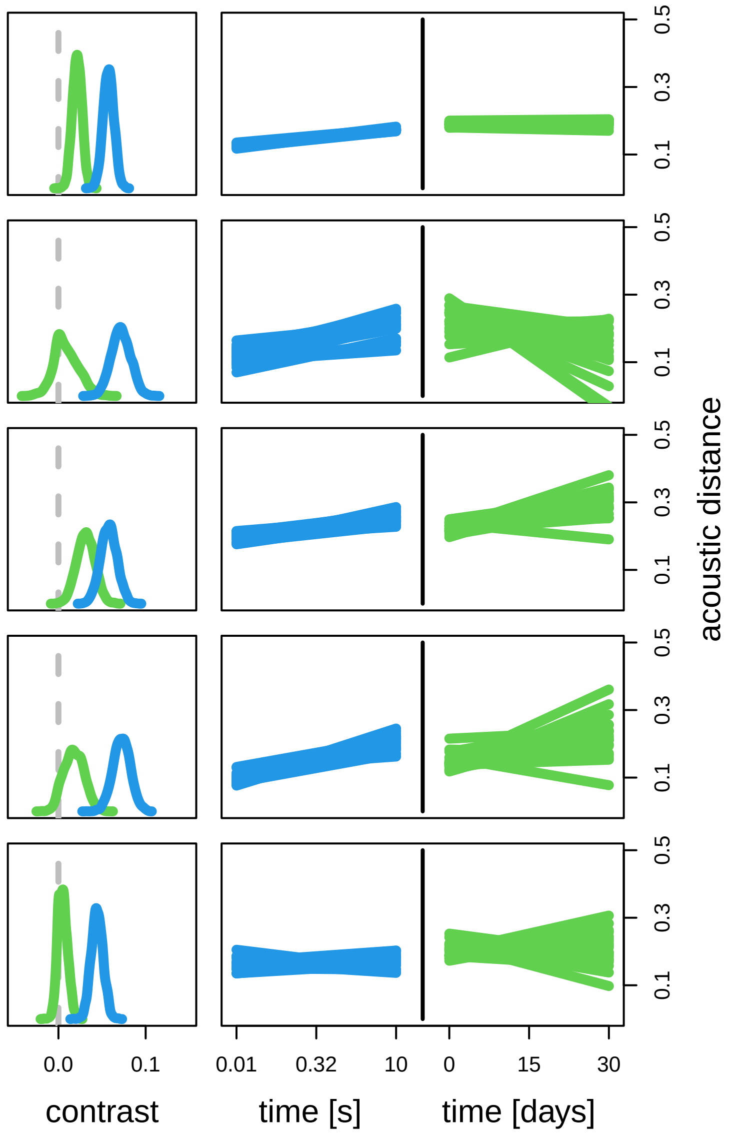

Figure S6: Model results for contact calls, tja calls, trruup calls, alarm calls and growls (ttb) using mel frequency cepstral coefficient cross correlation. Blue density plots are the posterior contrast between the similarity of calls from different individuals versus the same individual and same recording. Green density plots are the posterior contrast between the similarity of calls from different individuals vs from the same individual but different recordings. Blue lines are 20 samples from the posterior prediction of acoustic distance throughout time within a recording. Green lines are 20 samples from the posterior prediction of acoustic distance throughout days between a recording.

#### 2.4 pDFA

For the results presented in the paper there was no overlap with zero for any of the pDFAs. For full distributions and sample sizes see Figure S7.

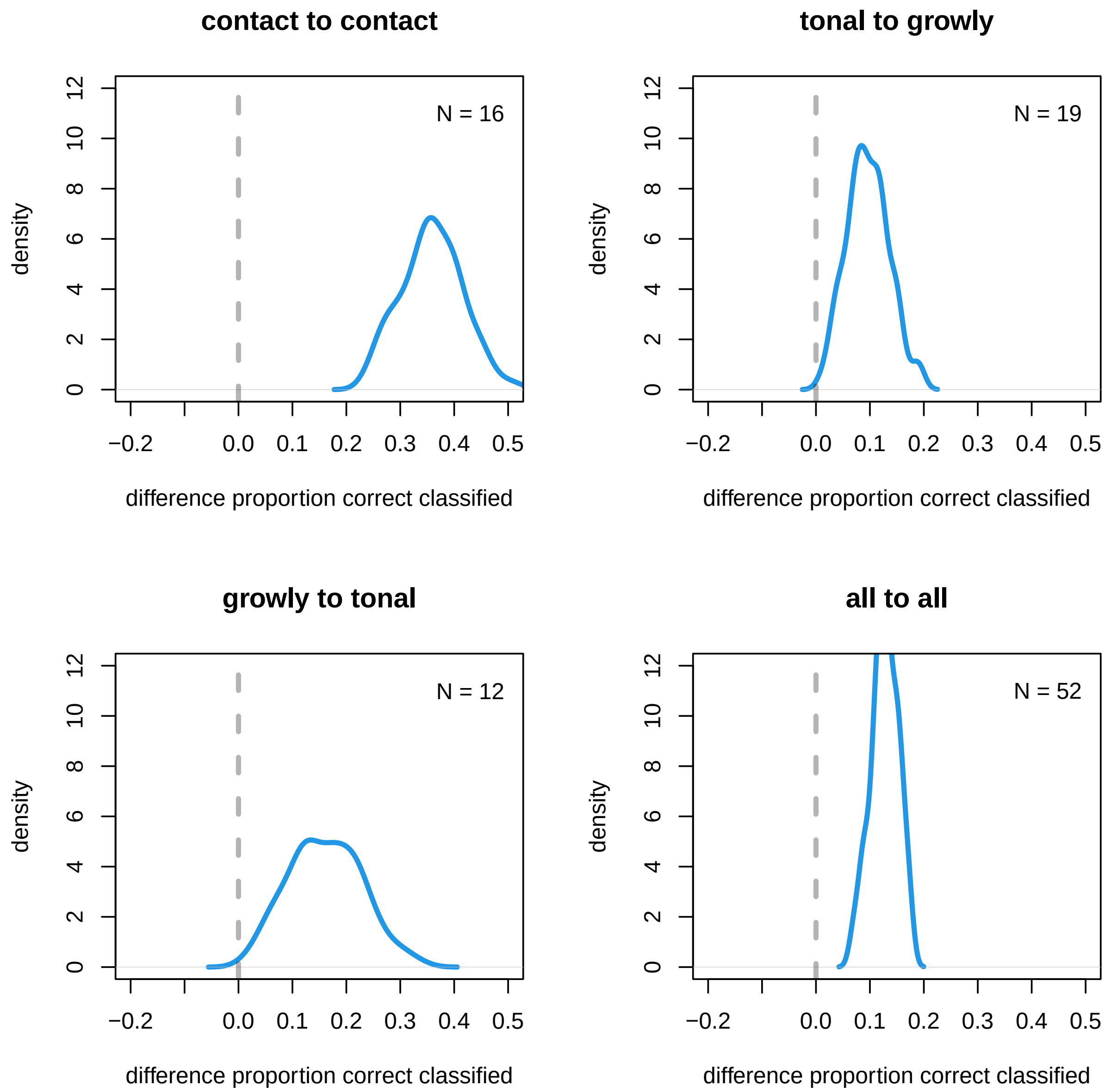

Figure S7: pDFA results. Densities of the differences between the trained and random DFAs. N is number of individals for which there was enough data available.

When sub-setting the data to only include females from Promenade Passeig de Lluís Companys the results for growly to tonal fell towards chance-level, but all other results remained stable with overlap with zero less than 2%. For full distributions and sample sizes see Figure S8.

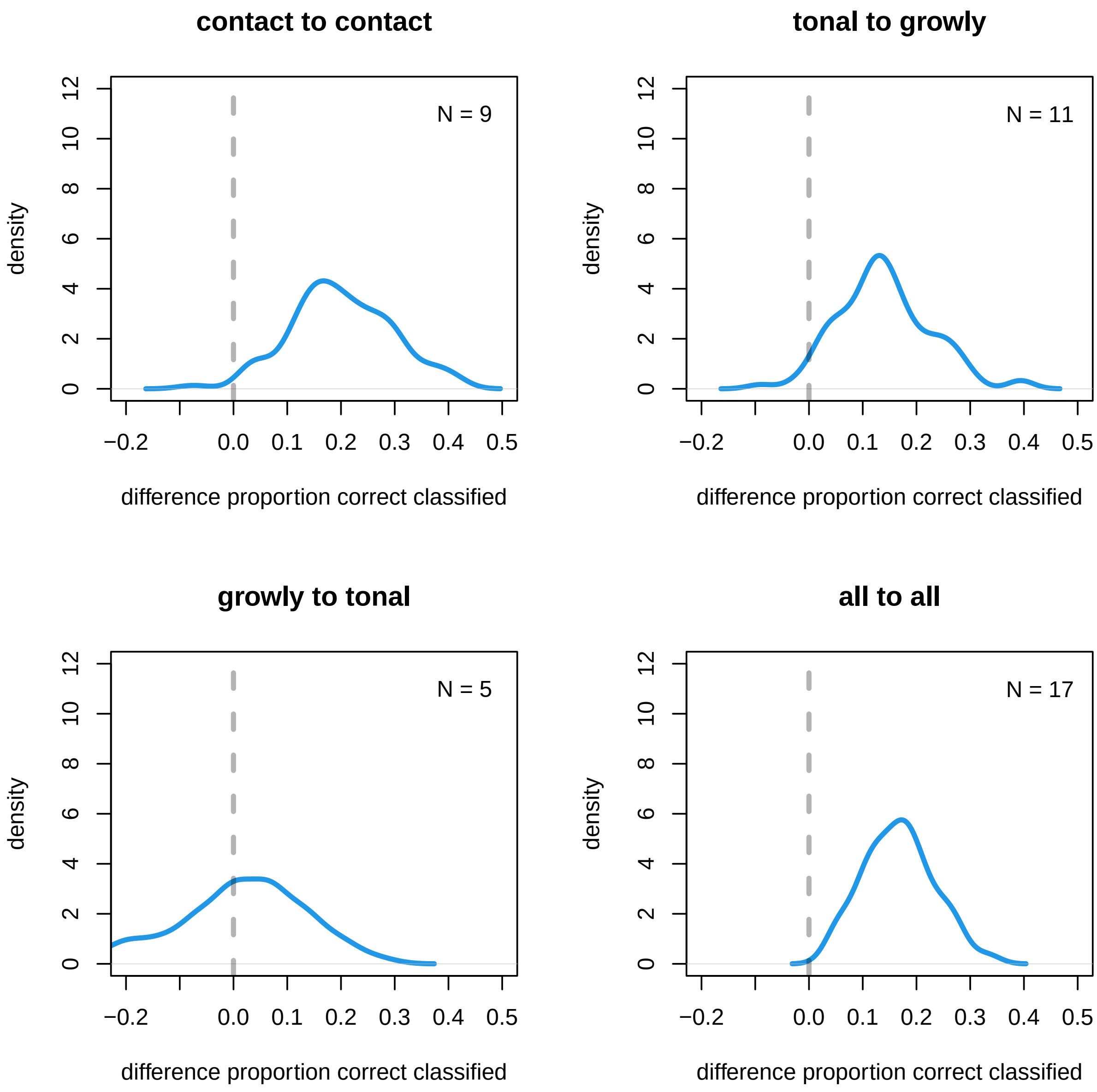

Figure S8: pDFA results for females from Promenade Passeig de Lluís Companys. Densities of the differences between the trained and random DFAs. N is number of individuals for which there was enough data available.

When sub-setting the data to only include males from Promenade Passeig de Lluís Companys the results for both cross-call type models fell towards chance-level. For full distributions and sample sizes see Figure S9.

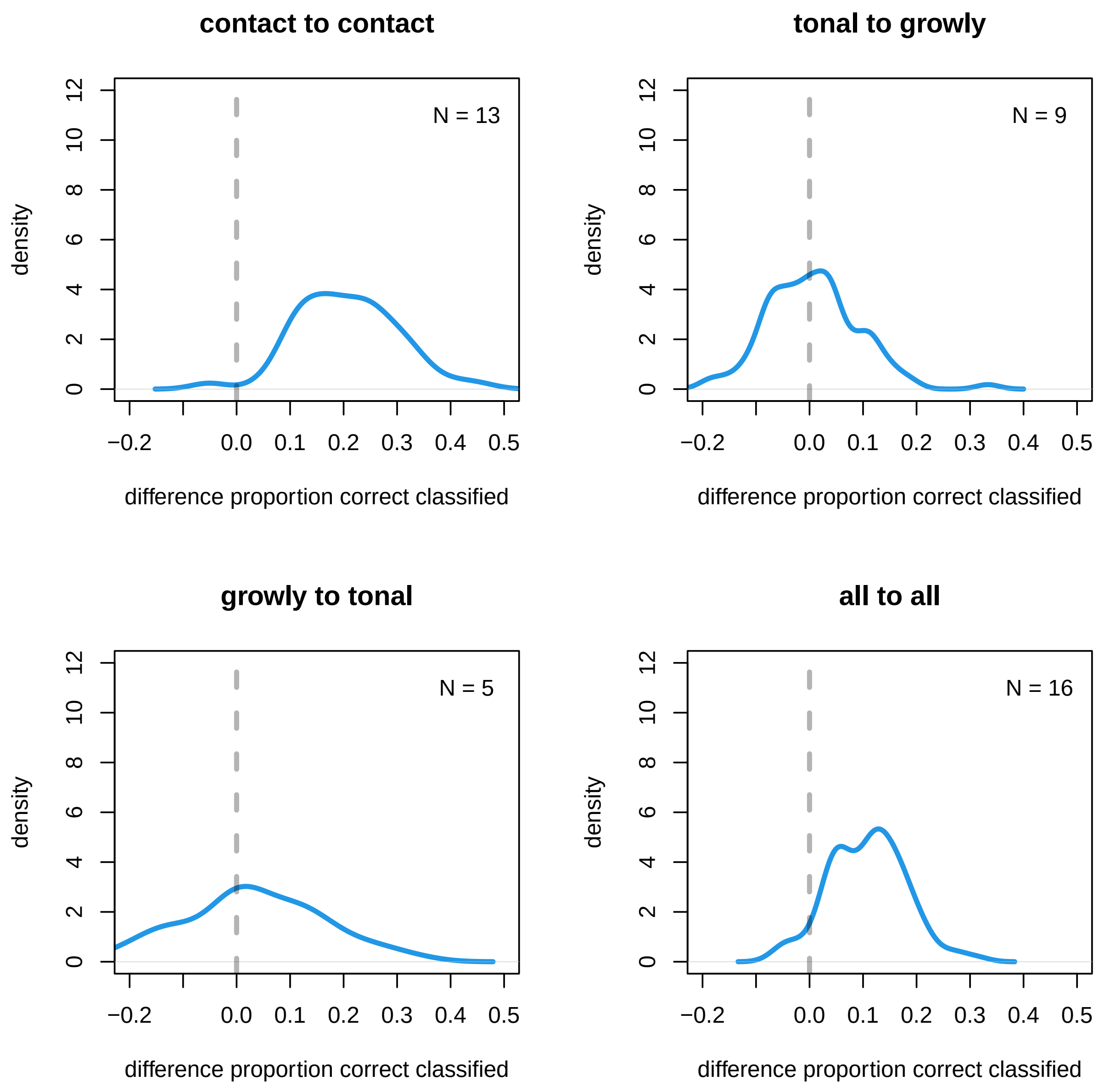

Figure S9: pDFA results for males from Promenade Passeig de Lluís Companys. Densities of the differences between the trained and random DFAs. N is number of individuals for which there was enough data available.

When randomising within Promenade Passeig de Lluís Companys and Parc de la Ciutadella the results for the cross call type models remained stable but overlap with zero increased to 6-7%. For full distributions and sample sizes see Figure S10.

Figure S10 and Figure S10 show the confusion matrices for one run of discriminant function analysis for the cross call type models.

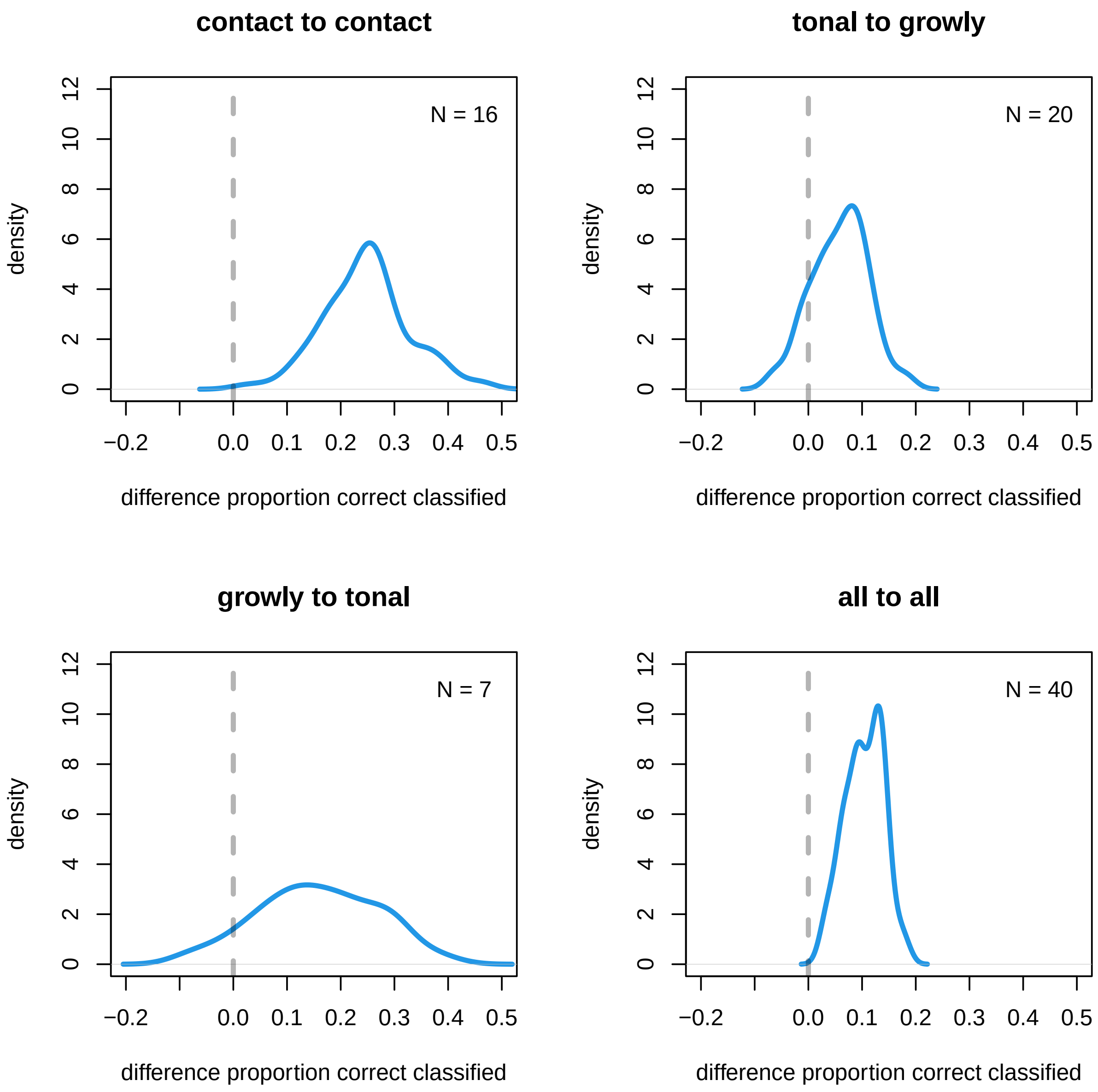

Figure S10: pDFA results with randomisation withing Promenade Passeig de Lluís Companys and Parc de la Ciutadella only. Densities of the differences between the trained and random DFAs. N is number of individals for which there was enough data available.

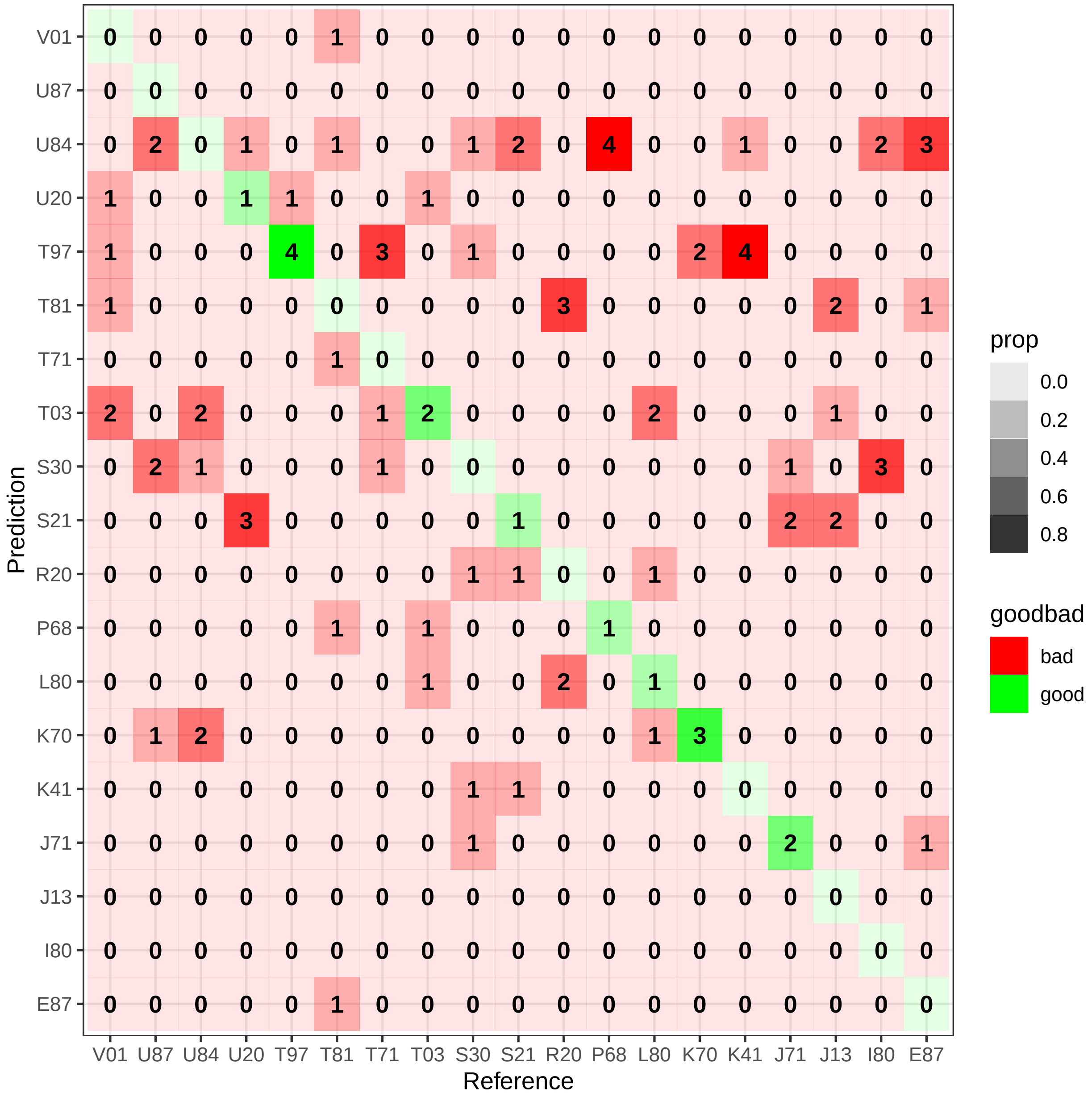

Figure S11: Confusion matrix for one run of discriminant function analysis where the model was trained on tonal calls and tested on growly calls. Numbers indicate how many calls were classified in that category, colour represent relative proportions.

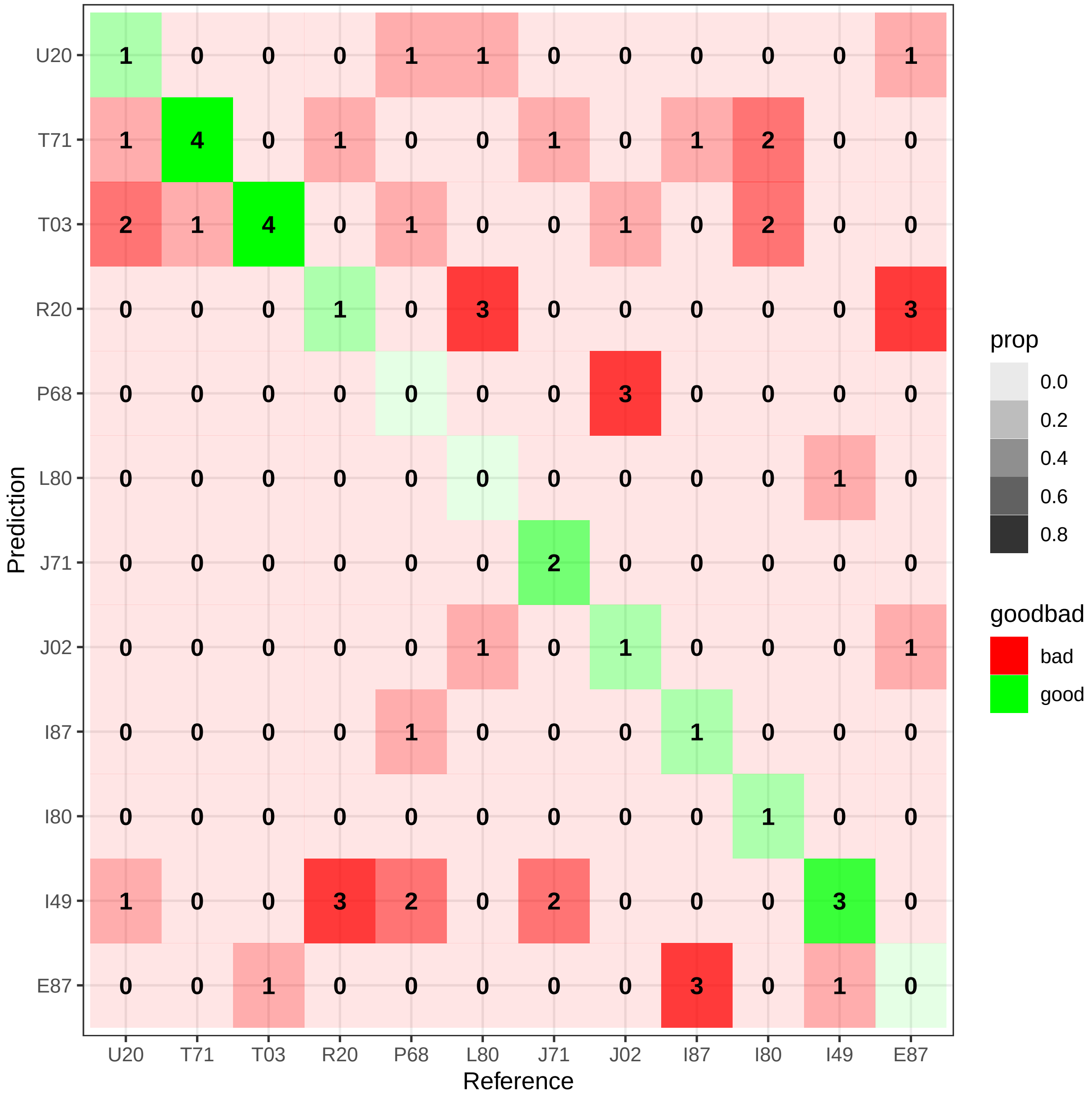

Figure S12: Confusion matrix for one run of discriminant function analysis where the model was trained on growly calls and tested on tonal calls. Numbers indicate how many calls were classified in that category, colour represent relative proportions.

#### 2.5 PCA - SPCC - individual signal

We ran principle component analysis on the distance matrix obtained from running SPCC. We subsetted the data to five random individuals to be able to visualise individuals with colours. Figure S13 shows some clustering in all call types but the *trruup* call. Note that multiple calls per recording are included, so that some of the clustering is due to recording.

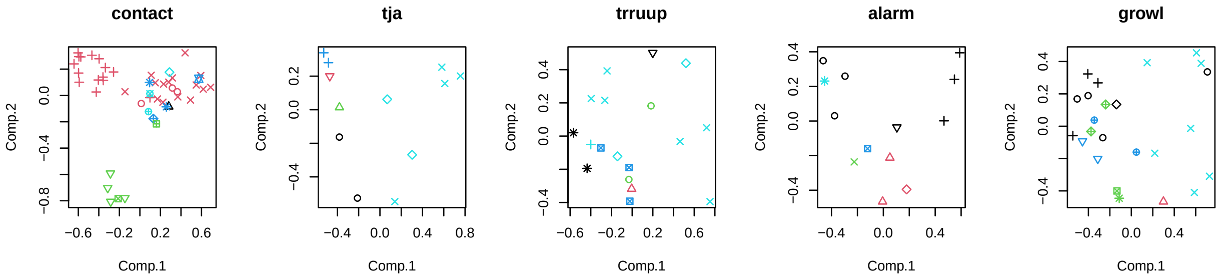

Figure S13: PCA scores for SPCC distance matrix of the main five call types. Colours represent individual. Shapes represent different recordings within individual.

Gabry, Jonah, and Rok Češnovar. 2021. *Cmdstanr: R Interface to 'CmdStan'*.

Gelman, Andrew, Daniel Lee, and Jiqiang Guo. 2015. "Stan: A Probabilistic Programming Language for Bayesian Inference and Optimization." *Journal of Educational and Behavioral Statistics* 40 (5): 530–43.

Giorgino, Toni. 2009. "Computing and Visualizing Dynamic Time Warping Alignments in r: The Dtw Package." *Journal of Statistical Software* 31: 1–24.

Kuhn, and Max. 2008. "Building Predictive Models in r Using the Caret Package." *Journal of Statistical Software* 28 (5): 1–26. https://doi.org/10.18637/jss.v028.i05.

Lachlan, RF. 2007. "Luscinia: A Bioacoustics Analysis Computer Program." *See Luscinia. Sourceforge. Net.[Google Scholar]*.

Ligges, Uwe, Sebastian Krey, Olaf Mersmann, and Sarah Schnackenberg. 2018. *tuneR: Analysis of Music and Speech*. https://CRAN.R-project.org/package=tuneR.

R Core Team. 2021. *R: A Language and Environment for Statistical Computing*. Vienna, Austria: R Foundation for Statistical Computing. https://www.R-project.org/.

Sueur, J., T. Aubin, and C. Simonis. 2008. "Seewave: A Free Modular Tool for Sound Analysis and Synthesis." *Bioacoustics* 18: 213–26. https://www.tandfonline.com/doi/abs/10.1080/09524622.2008.9753600.

Venables, W. N., and B. D. Ripley. 2002. *Modern Applied Statistics with s*. Fourth. New York: Springer. https://www.stats.ox.ac.uk/pub/MASS4/.
